# Multi-omics analysis reveals signatures of selection and loci associated with complex traits in pigs

**DOI:** 10.1101/2023.09.19.558553

**Authors:** Guoqiang Yi, Lei Liu, Yilong Yao, Yuwen Liu, Jiang Li, Yalan Yang, Lingzhao Fang, Delin Mo, Longchao Zhang, Yonggang Liu, Yongchao Niu, Liyuan Wang, Xiaolu Qu, Zhangyuan Pan, Lei Wang, Muya Chen, Xinhao Fan, Yun Chen, Yongsheng Zhang, Xingzheng Li, Zhen Wang, Yijie Tang, Hetian Huang, Pengxiang Yuan, Yuying Liao, Xinjian Li, Zongjun Yin, Di Liu, Dongjie Zhang, Quanyong Zhou, Wangjun Wu, Jicai Jiang, Yahui Gao, George E. Liu, Lixian Wang, Yaosheng Chen, Martien A M Groenen, Zhonglin Tang

## Abstract

Selection signatures that contribute to phenotypic diversity, especially morphogenesis in pigs, remain to be further elucidated. To reveal the regulatory role of genetic variations in phenotypic differences between Eastern and Western pig breeds, we performed a systematic analysis based on seven high-quality *de novo* assembled genomes, 1,081 resequencing data representing 78 domestic breeds, 162 methylomes, and 162 transcriptomes of skeletal muscle from Tongcheng (Eastern) and Landrace (Western) pigs at 27 developmental stages. Selective sweep uncovers different genetic architectures behind divergent selection directions for the Eastern and Western breeds. Notably, two loci showed functional alterations by almost fixed missense mutations. By integrating time-course transcriptome and methylome, we revealed differences in developmental timing during myogenesis between Eastern and Western breeds. Genetic variants under artificial selection have critical regulatory effects on progression patterns of heterochronic genes like *GHSR* and *BDH1*, by the interaction of local DNA methylation status, particularly during embryonic development. Altogether, our work not only provides valuable resources for understanding pig complex traits, but also contributes to human biomedical research.

## Introduction

Pigs (*Sus scrofa*), a major farm animal species worldwide, were domesticated independently in Anatolia and South China around ∼10,000 years ago^1^. Subsequent long-term natural and artificial selection contributed to the formation of many breeds with diverse phenotypic characteristics and environmental adaptations. Particularly, some Western breeds have been intensively selected over the past 100 years and show remarkable differences in economic traits like body weight, growth rate, and intramuscular fat percentage, as compared with Eastern breeds^2,3^. Besides providing an abundance of agricultural products, pigs can also serve as a valuable biomedical model for human diseases and are promising as potential organ donors for xenotransplantation^4-6^. Therefore, exploring the genetic changes and selection regimes behind pig domestication and breeding will have great value at biological, medical, and economic levels.

In the past decade, several studies have attempted to mine the significant consequences of pig domestication and suggested numerous quantitative trait loci (QTLs) and genes with potential functional implications in phenotypic diversification, like *ESR1*, *KIT*, *LCORL*, *MC1R*, *NR6A1*, and *PLAG1*^1,7-10^. Identifying these pivotal genes allows for accelerating the pig breeding process, whereas many more genetic determinants are still waiting to be discovered. Hence, a comprehensive genetic survey on larger sample sizes and more pig breeds is urgently requred. Furthermore, an increasing body of evidence has revealed much more prominent roles of regulatory elements in the domestication process due to a lack of detrimental pleiotropic effects as compared with coding mutations^11,12^. Variants in regulatory elements can affect the spatiotemporal expression patterns of key developmental genes, and transcriptional changes even subtle can influence the developmental timing of certain organs and might ultimately lead to phenotypic alterations^13^. Nevertheless, how artificial selection sculpts phenotypic diversity by reshaping the genome, transcriptome, and epigenome remains largely unknown.

To enhance our understanding of (epi)genetic substrates underpinning economic traits and especially their contributions to morphogenesis in pigs, we conducted a joint analysis of whole-genome resequencing data from 1,081 individuals, and time-course transcriptome and methylome data of skeletal muscle from 27 developmental stages in two representative Eurasian pig breeds. In this work, we established the most comprehensive pan-genome and genetic variation repertoire to date. We reveal plentiful selective sweep signatures underlying traits of economic values, and further illustrate many promising heterochronic genes and DNA methylation variations under artificial selection linked to myogenesis and meat production. Overall, these results provide novel insights into the (epi)genetic and phenotypic divergence between Eastern and Western pigs and nominate some promising determinants for future pig breeding programs and human biomedical research.

## Results

### A pan-genome sequence and genetic variation dataset in pigs

To capture a more comprehensive set of genomic sequences present in pigs than the reference genome Sscrofa11.1, we first performed PacBio High-Fidelity (HiFi) sequencing on one Asian wild boar (AW), three breeds from China (BMA: Bama Xiang, TT: Tibetan, and LT: Lantang) and three breeds from Europe (LW: Large White, BKS: Berkshire, and PTR: Piétrain) (Fig. 1a and Supplementary Fig. 1). An average coverage of 52.8× highly accurate long reads with per-base accuracy >99.9% per sample was obtained (Supplementary Table 1). Subsequently, we generated seven chromosome-scale genomes with an average length of 2.68 Gb by high-throughput chromosome conformation capture (Hi-C) based assembly (Supplementary Tables 2-3), which is slightly bigger than Sscrofa11.1 (2.68 Gb vs. 2.50 Gb). Transposable elements (TEs) makes up 38.8% of each genome ranging from 35.8% to 40.5% (Supplementary Table 4). The seven genomes have greater completeness and consistency based on the resulting assembly statistics and pair-wise colinear patterns compared to the reference genome from Duroc pigs. These genomes will serve as a rich resource for future research in pig genetics and genomics (Fig. 1b-c, Supplementary Figures 2-3 and Supplementary Tables 5-6). We further integrated two chromosome-level genomes, the Sscrofa11.1 reference, and a previous Luchuan genome reported by our group^14^, to build a high-resolution pan-genome (Supplementary Table 6). A total of 134.24 Mb non-redundant non-reference sequence with an N50 of 287,722 bp was obtained (Supplementary Table 7). We observed a higher GC content in the novel sequences than in the reference genome (49.5% vs. 41.4%) (Fig. 1d). Approximately 83.72% of the non-redundant non-reference sequences comprised repetitive elements, which is higher than the reference genome. In total, 1,099 novel genes were detected with a coding sequence (CDS) N50 size of 774. Of these, 94.54% (1,039 genes) were supported by transcript evidence, and 77.53% (852 genes) were successfully annotated for ≥1 function term (Supplementary Table 8). The Sscrofa11.1 assembly together with these putative pan-sequences were used as the final reference genomes for subsequent analyses.

**Fig. 1.**
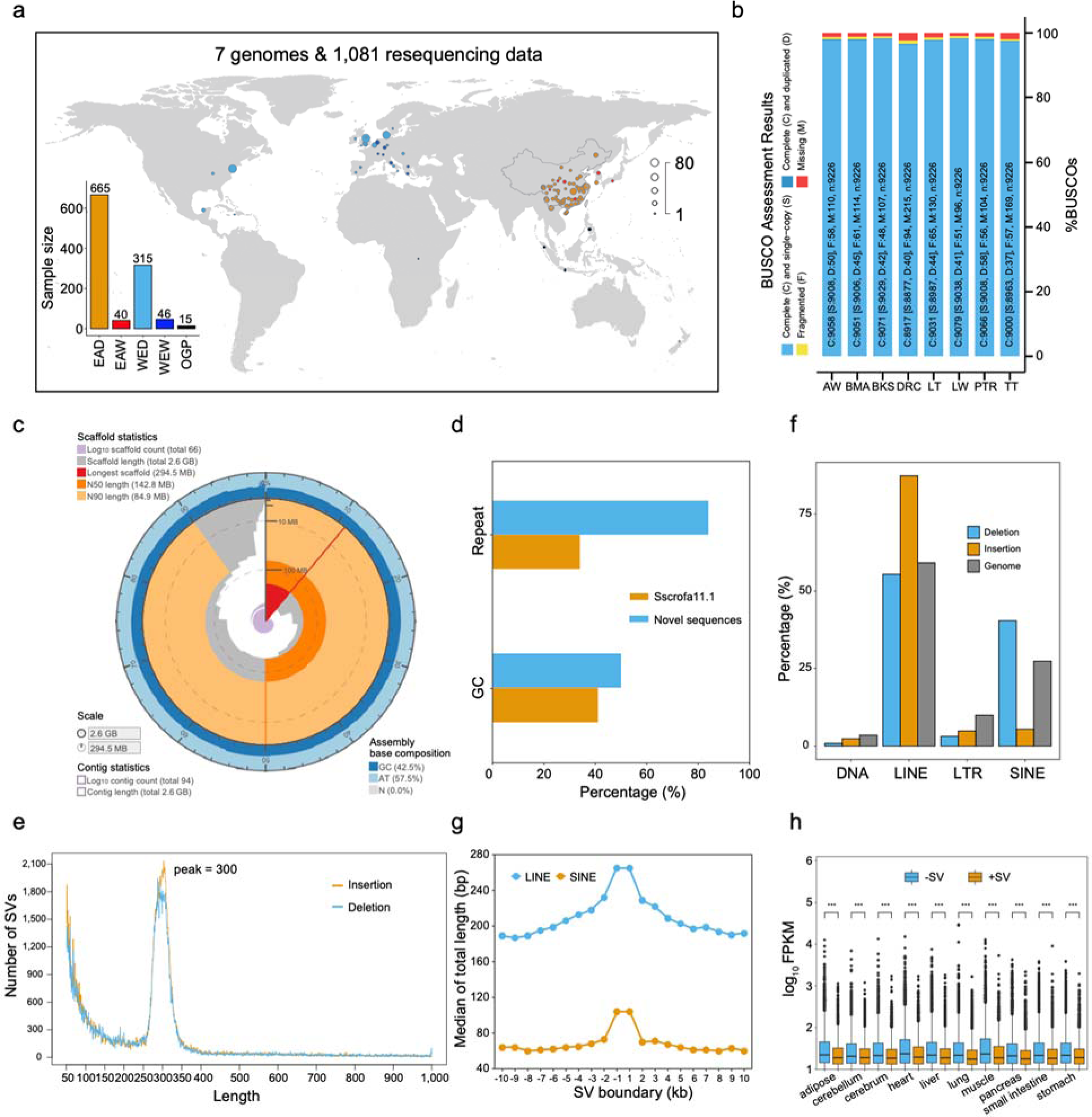
Comprehensive landscape of pig pan-genome. **a**, Geographic locations of all breeds and species for the five categories. The histogram shows the sample size of each category. EAD, Eastern domestic pigs; WED, Western domestic pigs; EAW, Eastern wild boars; WEW, Western wild boars; OGP, Outgroup (other Suids). **b**, BUSCO assessment results for genome assemblies of eight pig breeds. The Duroc genome (DRC) is the reference assembly which was downloaded from the Ensembl database. AW: Asian wild boar; BMA: Bama Xiang, BKS: Berkshire; LT: Lantang; LW: Large White; PTR: Piétrain; TT: Tibetan. **c**, Snail plots describing the assembly statistics of the Piétrain breed. Other six genomes were displayed in Supplementary Figure 2. **d**, The percentages of repeat sequence and GC content in the reference genome and novel pan-sequences. **e**, Size distribution of putative structural variants. **f**, Enrichment of structural variants in repeat elements. **g**, Distribution of repeat elements around structural variants boundary. **h**, The expression difference between genes with and without structural variants in multiple tisses.”

To enable a more complete and equitable understanding of genomic diversity, we anchored these assembled genome sequences onto the reference genome using the anchorwave program^15^. A total of 187,927 structural variations (SVs) with a peak at a size of 300 bp were found (Fig. 1e), which was consistent with previous results^16,17^. Notably, we found that long interspersed nuclear elements (LINEs) and short interspersed nuclear elements (SINEs) are two major contributors driving structural variations, and mainly enriched within SVs and around SVs boundaries (Fig. 1f-g). In addition, 7,667,617 small insertions and deletions (indels, referring ≤50 bp in this work) were identified (Supplementary Fig. 4). In the sequence fragment distribution plot, the even number was higher than the adjacent odd number (Supplementary Fig. 4). By leveraging multi-tissue RNA-seq data, we found that SVs generally exerted negative effects on the expression levels of flanking genes (Fig. 1h).

To build a rich genetic variation repertoire in pigs, we gathered whole-genome sequencing data from 1,081 individuals (Supplementary Tables 9-10), representing the majority of geographically diverse breeds across the world (Fig. 1a). We grouped all these individuals into five categories: an Eastern domestic group (EAD) including 665 pigs from 53 Chinese breeds (∼ 60% of all Chinese pig breeds) and one Korean indigenous breed, a Western domestic group (WED) containing 315 pigs from 25 typical Western breeds, an Eastern wild group (EAW) comprising 40 wild boars from Asia, a Western wild group (WEW) including 46 European wild boars, and 15 individuals from five different wild pig species (*Suidae*) like *Sus barbatus*, *Sus cebifrons*, *Sus celebensis*, *Sus verrucosus*, and *Phacochoerus africanus* that served as an outgroup (OGP). The entire data amounts to 45.3 Tb, representing the most extensive genetic diversity in pigs so far (Supplementary Table 11). For short variant discovery, a final set of 30,143,962 single-nucleotide polymorphisms (SNPs) and 5,496,594 insertions/deletions (INDELs) after removal of low-quality variants was identified (Supplementary Fig. 5a-b), which, to our knowledge, currently is the most abundant repertoire of genetic variants. These variants showed a clear depletion within exons but high levels of enrichment within introns and intergenic regions (Supplementary Fig. 5c). Most of the SNPs and INDELs located in intergenic (59.52% and 59.08%) and intronic (37.55% and 38.47%) regions, whereas only 0.56% of SNPs and 0.10% of INDELs were present in coding sequences (Supplementary Table 12).

### Population structure and potential selection signatures

Population structure analysis according to the neighbor-joining method displayed a clear subdivision into five clades, despite the presence of ancestry admixture between certain EAD and WED breeds (Fig. 2a). This inferred relationship was broadly consistent with principal component analyses (PCA) patterns, exhibiting a apparent separation between Eastern and Western pigs (Fig. 2b). By selecting the top 1% of putative selective sweeps based on computed composite likelihood ratios (CLR), we identified around 227-Mb with significant selection footprints in EAD and WED groups, respectively. The permutation test revealed that selection signatures were more enriched in introns, intergenic regions, and TEs compared to promoters and exons (Fig. 2c).

**Fig. 2.**
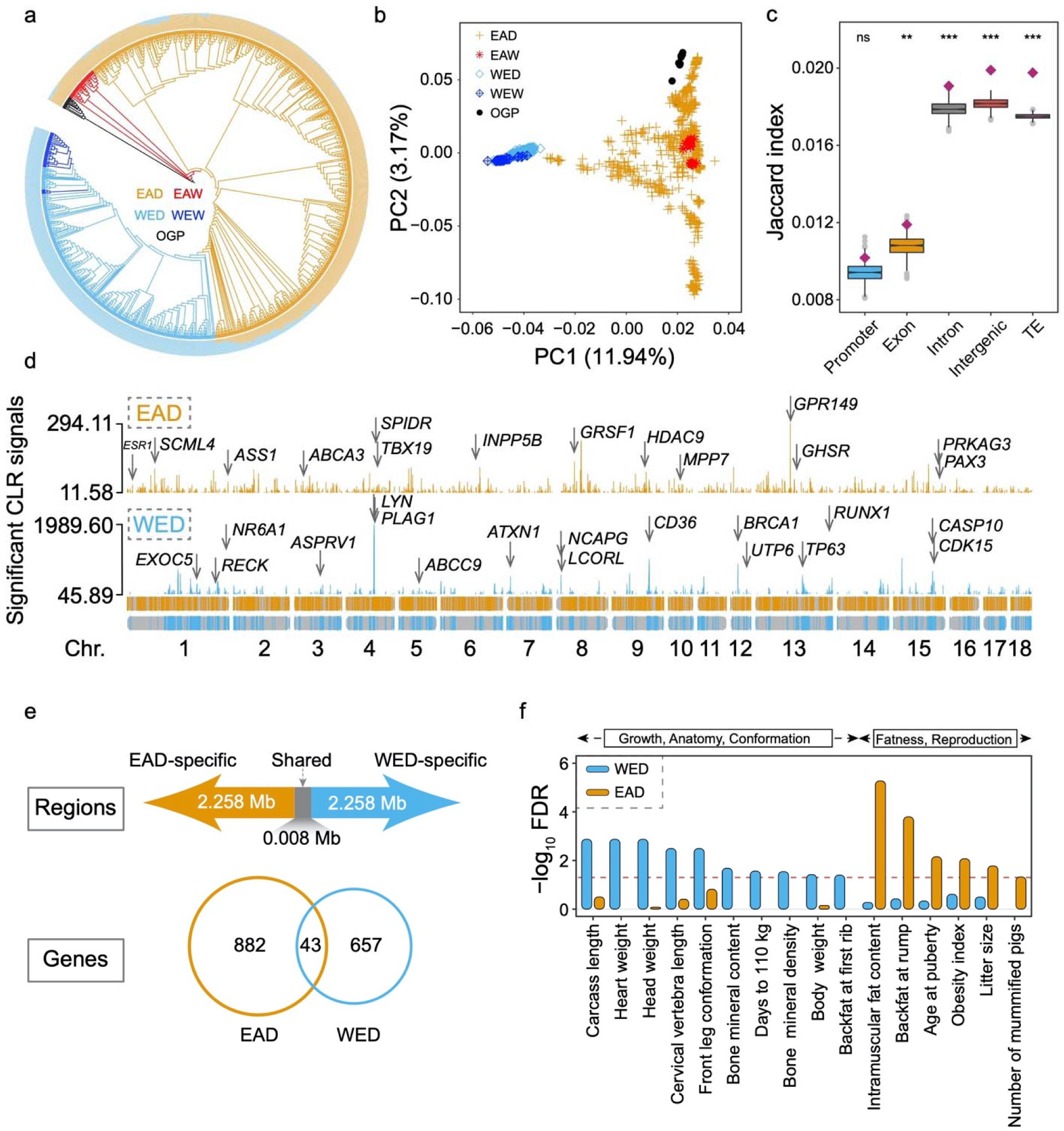
Population structure and selective sweeps in Eastern and Western pig populations. **a**, Population structure of all 1,081 individuals. The neighbour-joining phylogenetic tree includes 665 Eastern domestic pigs (EAD), 315 Western domestic pigs (WED), 40 Eastern wild boars (EAW), 40 Western wild boars (WEW), and 15 individuals from 5 other Sus species and *Phacochoerus africanus* (OGP). The circular barplot indicates the individual ancestry coefficient for each sample in which we set the K value as 2. **b**, Principal component analysis based on all putative autosomal SNPs. **c**, Overlap analysis between putative selective sweeps and promoter, exon, intron, intergenic and transposable elements by Jaccard index score. \**p* < 0.05; \*\**p* < 0.01; \*\*\**p* < 0.001; ns, not significant. **d**, Manhattan plots for all significant selective sweeps in Eastern and Western pig groups. Promising candidate genes are marked in the graph. This result implied that Western domestic pigs were subject to stronger selection pressures. **e**, The shared and unique genomic intervals and genes under selection between Eastern and Western pigs. **f**, Functional enrichment analysis of Eastern and Western swept genes against the Pig QTL Database.

Selective sweeps in WED showed higher signals and more concentrated locations (Fig. 2d) than those in the EAD group. We found 925 and 700 candidate genes in EAD- and WED-specific selective sweep regions (Supplementary Tables 13-14), respectively. Notably, the number of selection footprints and genes shared between Eastern and Western pigs was limited (Fig. 2e). Among all 1,625 putative genes under selection, we found many candidates reported by previous studies (Fig. 2d), including *CASP10*, *ESR1*, *LCORL*, *NCAPG*, *NR6A1*, and *PLAG1*^8-10,18^, suggesting high reliability of current predictions.

Besides, we provided a list of novel observations with respect to swept genes that were worth further exploration (Fig. 2d and Supplementary Fig. 6a), like *MPP7* for reproduction traits and *PRKAG3* for meat quality^18-20^. Many transcription factors (TFs) like *PAX3*, *RUNX1*, *TP63*, and *TBX19* involved in many vital biological processes were found to be under selection. Several swept intervals were located far from genes of particular functions like *E2F2* and *MYOG* and might act as intergenic or distal functional elements. Gene ontology (GO) enrichment analysis revealed that the swept gene set from EAD was significantly enriched in developmental growth and response to growth factor terms. In contrast, over-represented terms in the WED group mainly referred to the Wnt signaling pathway, cardiovascular system development and organelle localization (Supplementary Fig. 6b). We further probed the phenotypic consequences of swept genes by integration with the pig QTL database^21^ and provided more support for notably different selection directions between EAD and WED breeds. The hypergeometric test showed that traits driven by EAD swept genes were involved in fatness and reproduction classes, and genes under selection in WED group primarily concerned growth, anatomy, and conformation categories (Fig. 2f).

### Functional implication of coding variants linked to selective sweeps

Given that mutations within coding sequences might affect protein structure and function, we first explored the functional implications of these genomic variants. In total, we detected 156,654 autosomal coding SNPs at a genome-wide scale. By pruning SNPs outside the swept genes and SNPs with delta allele frequency (ΔAF) between EAD and WED groups ≤ 0.7, we finally kept 990 nearly fixed coding variants (Fig. 3a). The Ensembl Variant Effect Predictor (VEP) suite returned 626 SNPs (63.23%) to be synonymous, 339 (34.24%) as missense variants, 19 as splicing mutations, and six as stop-gain and start-loss mutations (Supplementary Table 15). Previous studies proposed that a missense substitution (chr 1_265347265_A/G, c.T575C or p.Pro192Leu) in the exon 5 of the *NR6A1* gene was the causative site affecting the number of vertebrae^8,22^. Consistently, we provide compelling evidence for this finding and illustrate the conservation and function of the amino acid substitution (Supplementary Fig. 7a-f).

**Fig. 3.**
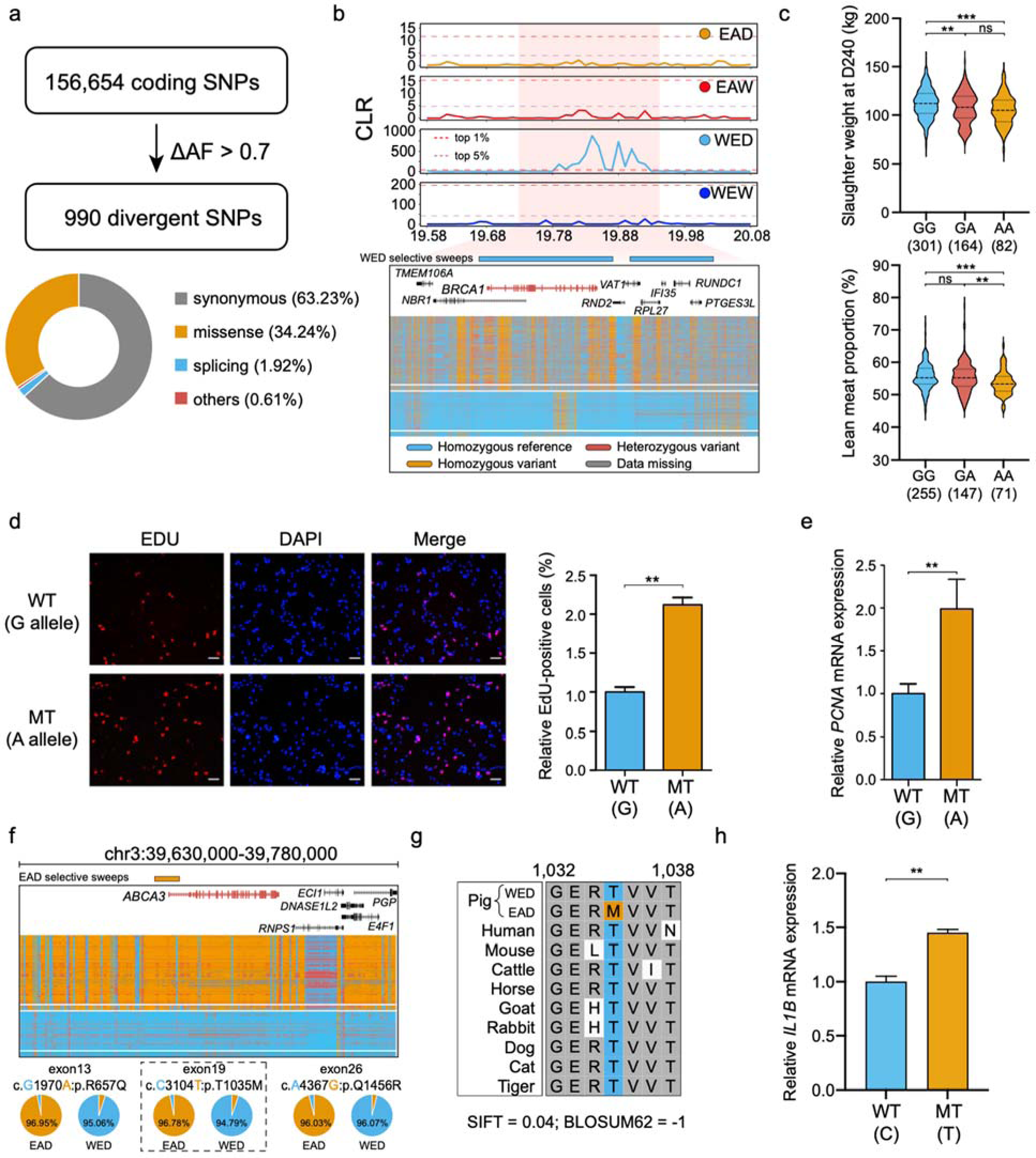
Functional implications of coding variants driven by selective sweeps. **a,** Annotation of coding variants with delta allele frequency (ΔAF) between EAD and WED groups larger than 0.7. **b**, The composite likelihood ratio (CLR) values and genotype patterns in the *BRCA1* region among EAD, EAW, WED and WEW groups. Dashed purple and red lines represent thresholds of the top 5% and 1% selective sweeps, respectively. **c**, Significant differences in slaughter weight at day 240 and lean meat proportion contributed by a coding SNP of the *BRCA1* gene. \**p* < 0.05; \*\**p* < 0.01; \*\*\**p* < 0.001; ns, not significant. **d**, Comparison of EdU proliferation assays for porcine adipocyte between the wild-type and mutant alleles. **e**, Increased proliferation capacity of mutant allele according to higher expression of *PCNA* marker by qRT-PCR. **f**, The CLR values and genotype landscape in the *ABCA3* region among EAD, EAW, WED and WEW groups. Dashed purple and red lines represent thresholds of the top 5% and 1% selective sweeps, respectively. Three nearly fixed coding variants were found in this region. **g**, Multiple sequence alignment of amino acids across ten representative mammalian species. **h**, Alleviated immune injury of wild-type allele according to lower expression of *IL1B* marker by qRT-PCR.

**Fig. 4.**
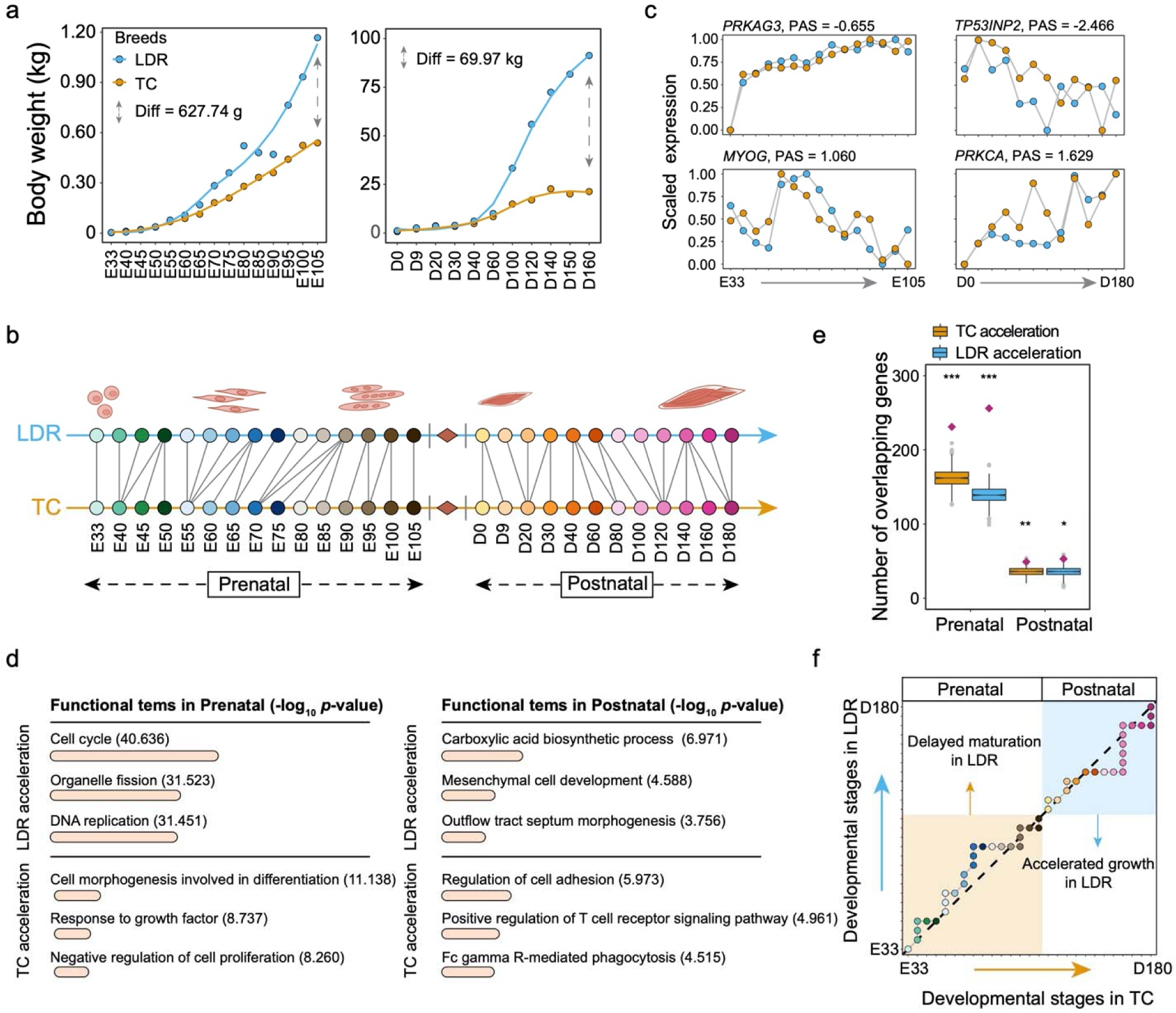
Inter-breed heterochrony of porcine skeletal muscle development by swept genes. **a**, Differences in body weight of Landrace (LDR) and Tongcheng (TC) pigs at multiple developmental stages. **b**, Cross-breed correspondences of skeletal muscle developmental stages in LDR and TC pigs. **c,** Examples of differentially progressing genes in the prenatal and postnatal stages. Based on the definition of TimeMeter, positive PAS indicates a faster expression pattern in TC (TC acceleration genes), while negative means accelerated changes in LDR (LDR acceleration genes). PAS, progression advance score. **d**, Gene ontology enrichment analysis predicted with LDR and TC acceleration genes in prenatal and postnatal stages. \**p* < 0.05; \*\**p* < 0.01; \*\*\**p* < 0.001; ns, not significant. **e**, Significant overlaps between differentially progressing genes (DPGs) and swept genes. The DPG set was predicted from time-course RNA-seq data from LDR and TC pigs. Y-axis is the observed (purple diamond) and expected (boxplot) number of overlaps determined by 1,000 permutation tests. Permutation *p*-value threshold = 0.05, \**p* < 0.05; \*\**p* < 0.01; \*\*\**p* < 0.001; ns, not significant. **f,** Developmental heterochrony analysis based on only swept genes of LDR and TC pigs. Using LDR as a reference, TimeMeter implements a dynamic time warping algorithm in all RNA-seq data, to select the best alignment between the time series based on stage transcriptome correlations. This finding manifested that the differences in skeletal muscle development between TC and LDR can be traced back to very early stages of myogenesis, in which LDR showed more advanced and prolonged timing for myoblast proliferation.

**Fig. 5.**
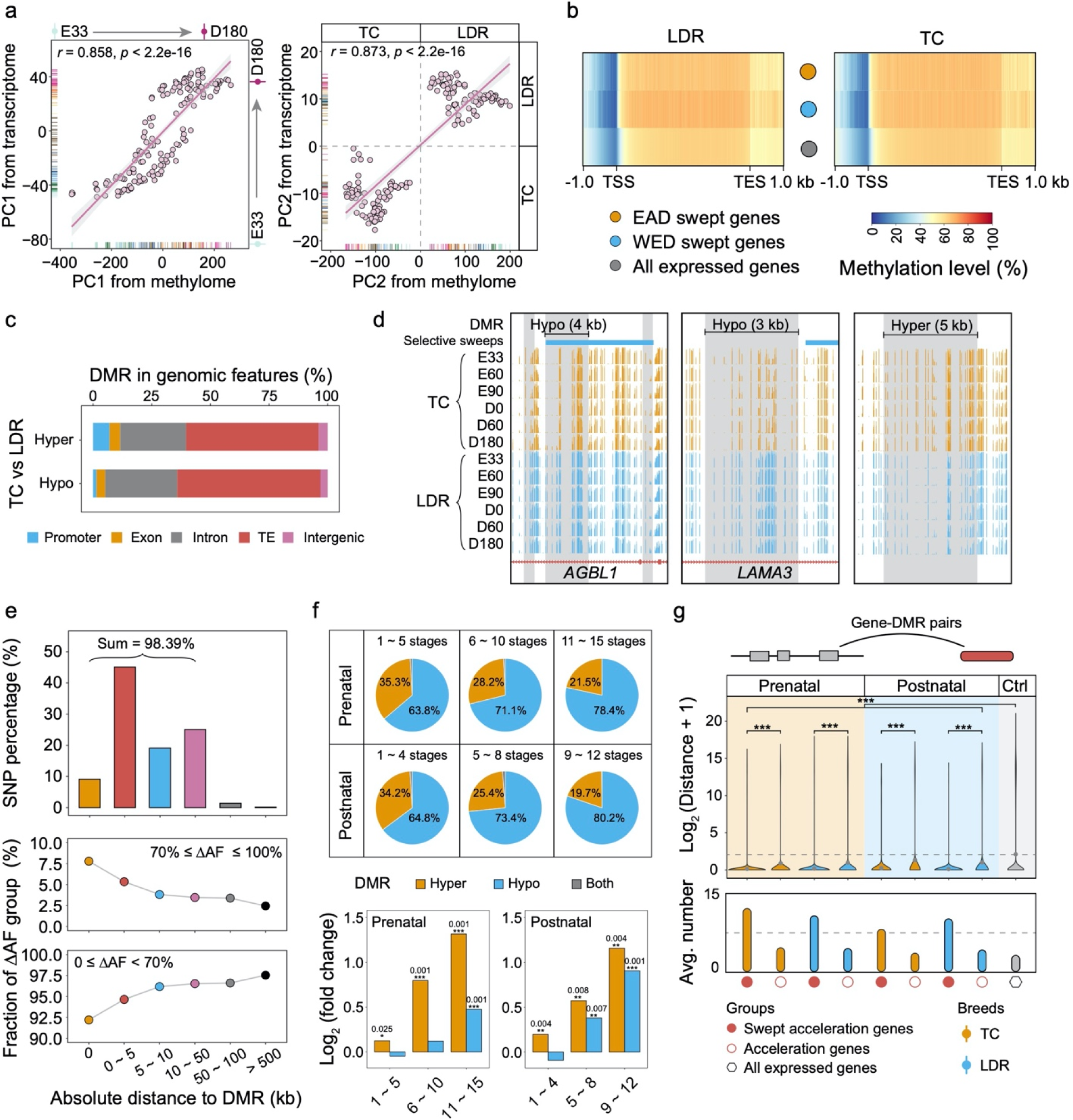
Differences in DNA methylome during pig domestication and breeding. **a**, Correlation analysis between methylome and transcriptome for PC1 and PC2. The PC1 and PC2 values were individually calculated from RNA-seq and WGBS data. **b,** Comparison of DNA methylation levels among EAD selective genes, WED selective genes and all expressed genes. **c**, The stacked graph of the proportion of DMRs in five different genomic features including promoter, exon, intron, TE and intergenic regions. **d**, Snapshot of UCSC genome browser showing five significant DMR loci colored by gray and selective sweeps in six representative stages. E, prenatal stages; D, postnatal stages. **e**, The absolute distance between DMR and the nearest SNPs at a genome-wide level. **f**, Relationships between DNA methylation variation and selective sweeps during prenatal and postnatal skeletal muscle development. \**p* < 0.05; \*\**p* < 0.01; \*\*\**p* < 0.001; ns, not significant. **g**, Gene-DMR pairs analysis for three gene categories. The distance and number of DMRs assigned to each group were calculated by BEDTools *closest* option.

**Fig. 6.**
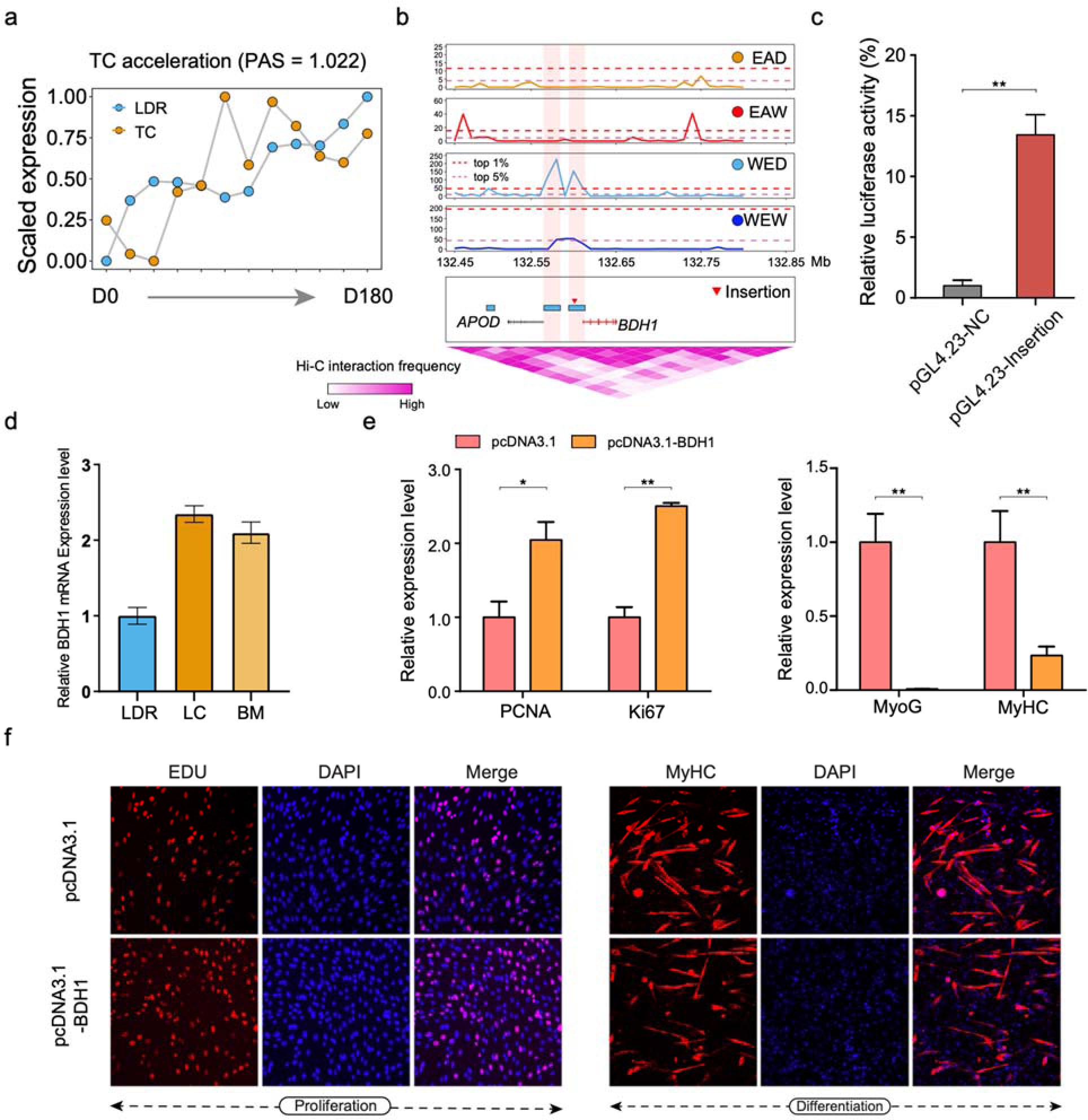
Comprehensive analysis of the *BDH1* gene related to skeletal muscle development. **a**, TimeMeter analysis showing the accelerated expression of *BDH1* in the TC postnatal stage. **b**, Selective sweeps regions in the four groups. Dashed purple and red lines represent thresholds of the top 5% and 1% CLR values in the *BDH1* region, respectively. **c**, Luciferase reporter assays in HEK293T cells to compare enhancer activity between the insertion. \**p* < 0.05; \*\**p* < 0.01; \*\*\**p* < 0.001; ns, not significant. **d**, Higher gene expression of *BDH1* in Chinese native pig breeds by qRT-PCR. **e**, Cell proliferation assessment for the expression level of *PCNA*, *Ki67*, *MyoG*, and *MyHC* marker upon *BDH1 overexpression* by qRT-PCR analysis in C2C12 cells. **f**, Proliferation and differentiation assays of murine C2C12 cells following *BDH1* overexpression.

Besides the *NR6A1* gene, we also highlighted several promising candidates related to traits of economic importance. We found two broad locations with a strikingly high signal of selective sweeps only in the WED group (Fig. 3b), which harbored eight functional genes including *BRCA1*, *NBR1*, and *RPL27*. Notably, further inspection of this region uncovered a coding mutation of c.G965A (chr12_19812845_G/A) located within exon 10 of the *BRCA1* gene. In this site, the mutant-type A allele was dominated in the EAD group (AF = 91.67%) but nearly absent in the WED group (AF = 3.27%) (Supplementary Fig. 8a-c). Meanwhile, this variant tends to be mildly conserved and possibly deleterious based on Ensembl SIFT (0.06) and BLOSUM62 (-1) scores as well as multi-species alignment (Supplementary Fig. 8d). To understand the effects of this variant, we compared the phenotypic differences of all three genotypes in 589 F2 individuals derived from the a cross between five Large White (European, WED) boars and sixteen Min (Chinese, EAD) sows^23^. We observed that this SNP has an allele substitution effect of 3.55 kg for slaughter weight at 240 days (SW) and 0.62% for lean meat proportion (LMP). At this locus, the AA genotype presented prominently lower SW and LMP than GG and GA genotypes (Fig. 3c). By *in vitro* adipocyte proliferation assays, the expression of G322D BRCA1 displayed enhanced capacity in adipocyte growth and development (Fig. 3d-e), corresponding to more significant fat deposition in obese EAD pigs, in contrast to WED breeds with the wild-type G allele.

Moreover, a swept region overlapping the *ABCA3* gene was of particular interest to us due to its highly distinct genotype distribution between EAD and WED groups and lung-specific expression of this gene (Fig. 3f and Supplementary Fig. 9a-b). We pinpointed three non-synonymous mutations displaying nearly complete LD among one another and prioritized the variant (chr3_39723445_C/T) in exon 19 as the top candidate, in consideration of its more deleterious effect on protein structure and function (SIFT = 0.04, BLOSUM62 = -1) (Fig. 3g and Supplementary Fig. 9c). Almost all individuals in the EAD group carried the alternative allele (T) at this site, whereas WED pigs were nearly fixed for the C allele. Since *ABCA3* mutations can cause surfactant deficiency, alveolar cell injury, and inflammation^24,25^, we hypothesized that the *ABCA3* gene was an intriguing genetic determinant responsible for breed-specific resistance/susceptibility against the pathogen Mycoplasma hyopneumoniae (Mhp)^25,26^. Experimental data via *in vitro* Mhp infection showed that the wild-type C allele was associated with lower expression levels of the *IL1B* gene (Fig. 3h), which indicated an alleviated immune response and lung injury for this allele.

### Swept genes regulated developmental heterochrony of skeletal muscle between EAD and WED groups

The enrichment analysis of swept genes against the pig QTL database illustrated large differences in growth and carcass characteristics between EAD and WED groups (Fig. 2f), intriguing us to compare the developmental trajectories of skeletal muscle across pig breeds. Thus, we designated Tongcheng (TC, an Eastern obese-type breed) and Landrace (LDR, a classical Western lean-type breed) pigs as research models. Phenotype records revealed that the two breeds showed high divergence in growth rate and muscle mass (Fig. 4a), and LDR pigs being more than twice as heavy as TC pigs at 160 days of age. Subsequently, we newly implemented a time-series RNA-seq experiment of 27 time points from skeletal muscle in TC, including 15 prenatal and 12 postnatal stages (Supplementary Table 16). Comprehensive analyses showed globally consistent transcriptional profiling across skeletal muscle development between TC and LDR^27^ (Supplementary Fig. 10a-f and 11a-d). We subsequently applied weighted gene co-expression network analysis (WGCNA) to explore critical functional modules implicated in skeletal muscle development. A total of 18 modules were found (Supplementary Fig. 12a), three of which showed significant overlap with putative swept genes and were primarily concerned with cell division, morphogenesis, and development processes (Supplementary Fig. 12b-c).

Based on these findings, we reasoned that the developmental heterochrony of skeletal muscle cells might be a vital impetus to divergent muscle mass. Therefore, we performed a comparative transcriptome analysis based on the dynamic time warping (DTW) algorithm in TimeMeter^28^ to dissect time shift patterns during skeletal muscle development between TC and LDR. We provided clues to mild heterochrony throughout skeletal myogenesis, featured by more advanced and prolonged myoblast proliferation in the prenatal stage and faster muscle growth in the postnatal period for LDR, whereas more accelerated myoblast maturation in the prenatal period for TC (Fig. 4b and Supplementary Fig. 13a). In total, we detected 4,947 and 1,184 differentially progressing genes (DPGs) in prenatal and postnatal stages, respectively, including *MYOG*, *TP53INP2*, and *MEF2B* (Fig. 4c, Supplementary Fig. 13b and Supplementary Tables 17-18). Based on calculated progression advanced scores (PAS), positive values refer to more advanced expression patterns in developmental progression in TC than those in LDR, which were defined as TC acceleration genes, while negative means accelerated changes in LDR (LDR acceleration genes). Functional enrichment analysis emphasized that LDR acceleration genes in the prenatal stage were significantly over-represented in cell cycle and DNA replication. In contrast, TC acceleration genes mainly contributed to cell differentiation and maturation (Fig. 4d). In the postnatal period, genes with more advanced progressions in LDR were persistently enriched for cell development and biosynthetic process, while TC advanced genes were linked to immune response.

To further elucidate the genetic basis underlying inter-breed differences in developmental timing, we evaluated the contributions of swept genes in tuning skeletal myogenesis. Compared with all genes in the pig genome, genes under selection showed higher expression levels at all stages for each breed (Supplementary Fig. 14), implying their essential functions in specific important biological processes. Subsequent permutation tests reported a highly significant overlap between DPGs and swept genes, especially in the prenatal phase (Fig. 4e). We conducted the same DTW analysis by focusing on these swept genes and uncovered highly consistent dynamic patterns with results from all expressed genes (Fig. 4f).

### Selective sweeps regulated gene expression through reshaping DNA methylation pattern

DNA methylation is one of the main epigenetic factors regulating gene expression in eukaryotes. To elucidate the contributions of DNA methylation variations in pig domestication and breeding, we performed whole-genome bisulfite sequencing (WGBS) in skeletal muscle across 27 developmental stages (Supplementary Table 19), corresponding to the aforementioned samples with RNA-seq data. We generated a global DNA methylome landscape in TC and LDR and found higher CpG methylation levels in TC (Supplementary Fig. 15 and 16a-c). Furthermore, WGBS-based PCA revealed concordant classification patterns with RNA-seq results (Supplementary Fig. 16d). Both the PC1 and PC2 between methylome and transcriptome showed strong curvilinear relationships, suggesting a clear classification of developmental trajectories and breeds (Fig. 5a). Functional correlations between these two types of datasets again confirm the results for promoter and gene body regions (Supplementary Fig. 17-20).

To understand the impacts of selective sweeps on DNA methylations, we performed a correlation analysis between WGBS and RNA-seq data based on only these swept genes. We found a reduced regulatory relationship in promoters, but enhanced patterns in gene bodies with average scores from zero to 0.250, compared with results using both all expressed genes and randomly sampled genes (Supplementary Fig. 21a-b). This finding highlights that sequence contexts under natural and artificial selection could reshape global expression patterns throughout the entire development due to DNA methylation variation. Compared with all expressed genes, promoter regions of putative swept genes generally have lower DNA methylation levels in both TC and LDR breeds, whereas gene bodies displayed overall higher levels of DNA methylation (Fig. 5b and Supplementary Fig. 22-23). By pair-wise differential methylation analyses, we obtained 166,265 differentially methylated regions (DMRs) domains (Supplementary Table 20), approximately 65% of which were hypomethylated in TC compared to LDR. The DMR intersection with five genomic features revealed that both hypermethylated and hypomethylated regions were primarily found in intron and TE regions (Fig. 5c). Visualization in the UCSC genome browser exemplified five significant DMR loci with distinct distribution patterns (Fig. 5d).

Based on the absolute distances between DMR domains and the nearest SNPs, we found that most DMR (98.39%) were located within 50 kb from identified SNPs, and around 55% had proximity within 5 kb (Fig. 5e). Furthermore, results indicated that these nearly fixed SNPs with ΔAF more than 0.7 were more prone to being in the vicinity of DMRs, as opposed to SNPs with ΔAF below 0.7. To further clarify the importance of DNA methylation in pig domestication and its relationship with genetic selection, we first separated DMR into three groups regarding the number of stages with significant differences in prenatal and postnatal stages, respectively. We found that the category with persistent DNA methylation variations at almost all stages (11 ∼ 15 stages in prenatal and 9 ∼ 12 stages in postnatal) possessed a higher proportion of Hypo-DMR domains than the other two groups (Fig. 5f, upper), supporting our results mentioned above. Enrichment analysis ascertained by permutation tests showed that DMRs with sustained changes were prone to being covered by selective sweeps in both prenatal and postnatal periods (Fig. 5f, lower). By mapping DMRs to the nearest genes, we found a much shorter distance and higher number of gene-DMR pairs for these acceleration genes under selection, compared with acceleration genes without selection and all expressed genes (Fig. 5g).

### Genetic basis of candidate genes in skeletal muscle development and meat performance

We first explored the regulatory roles of structural variations on candidate genes. An intriguing candidate was the *BDH1* gene, which was mainly involved in the regulation of the ketone metabolic process in skeletal muscle, liver, and heart tissues. The *BDH1* gene showed a more accelerated expression pattern in TC skeletal muscle at postnatal stage than in LDR (Fig. 6a), and we identified a 272 bp insertion located upstream of this gene in Eastern pig breeds. The insertion was covered by a selective sweep with strong signal (Fig. 6b). We speculated that this mutation exerted a positive regulatory role in the *BDH1* gene by a potential chromatin interaction, according to the 3D interaction heatmap (Fig. 6b). Luciferase reporter assays indicated increased enhancer activity of this insertion, and Chinese native pig breeds showed higher expression levels of *BDH1* than LDR breeds by qRT-PCR experiments (Fig. 6c-d). Further gene overexpression/suppression analysis revealed enhanced proliferation capacity but weak myoblast differentiation rates (Fig. 6e- f and Supplementary Fig. 24).

We next focused on the functional consequences of CpG-SNP mutations from methylated CpG to TpG/CpA in the putative selective sweeps and explored their regulatory functions on heterochronic genes and skeletal muscle development. Furthermore, we found more advanced progressions (PAS = -0.814) of the *GHSR* gene in LDR than TC during the embryonic period (Fig. 7a). This result suggested that *GHSR* should be a key contributor to accelerating myoblast proliferation. Multiple experiments supported that *GHSR* could promote myoblast proliferation but inhibit their differentiation and fusion *in vivo* and *in vitro* (Fig. 7b-c and Supplementary Fig. 25). Meanwhile, a pronounced selection signal was detected in the intron of *GHSR*, which harbored two significant Hypo-DMRs (Fig. 7d). Further analysis in this region revealed a CpG-SNP (chr13_111051076_C/T) almost perfectly fixed for the mutant allele in the EAD population and TC breed, which completely removed methylation status in TC due to the transition from C to T allele (Fig. 7e). We discovered that the SNP created a new transcription factor binding site of TBX21 via the alternate T allele by motif scan analysis. Subsequent luciferase reporter assays revealed higher enhancer activity for the T allele, which might be the critical contributor delaying the expression of *GHSR* in TC (Fig. 7f). The estimated allele substitution effect of this SNP for slaughter weight at D240 was 4.22 kg, and the individuals carrying one or two copies of the dominant T allele showed a significantly lower slaughter weight (Fig. 7g). Based on publicly available Hi-C data of porcine skeletal muscle, we revealed that the CpG-SNP was likely to regulate the expression of the *GHSR* gene since both are located in the same topologically associated domain (TAD) (Fig. 7h). Additionally, we also proposed that the heterochronic candidates, *CD36* and *NCAPG*-*LCORL* cluster^8,29-31^, were pivotal contributor to skeletal muscle development and slaughter weight in pigs based on similar strategies (Supplementary Fig. 25-28).

**Fig. 7.**
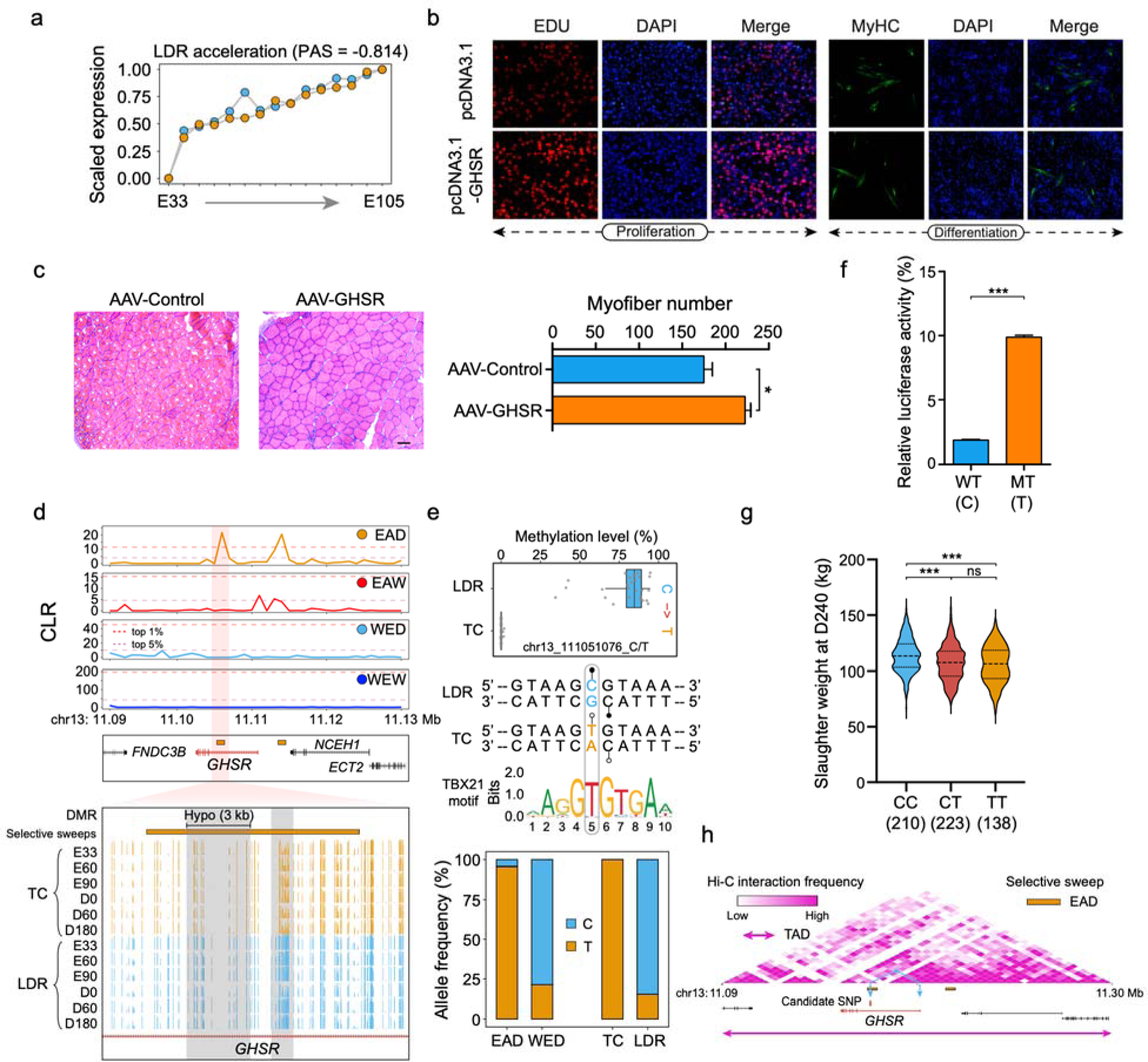
Comprehensive analysis of the *GHSR* gene related to skeletal muscle development and meat performance. **a**, TimeMeter analysis showing the accelerated expression of *GHSR* in the LDR prenatal stage. **b**, Proliferation and differentiation assays of murine C2C12 cells following *GHSR* overexpression. **c**, H&E staining and myofiber number of mice skeletal muscle at 10 days after *in vivo* injection of AAV-mediated non-target control and pcDNA3.1-GHSR, respectively. Representative images are shown at 20× magnification (scale bars = 100 μm). **d**, Selective sweeps reshaping DNA methylation levels across six representative developmental stages in LDR and TC pigs. Dashed purple and red lines represent thresholds of the top 5% and 1% CLR values in the *GHSR* region, respectively. **e**, The fixed C to T substitution in *GHSR* resulting in the removal of DNA methylation levels and the creation of a new transcription factor binding site of TBX21. **f**, Luciferase reporter assays in HEK293T cells to compare enhancer activity between the two alleles. \**p* < 0.05; \*\**p* < 0.01; \*\*\**p* < 0.001; ns, not significant. **g**, Higher slaughter weight at day 240 of CC genotype. **h**, The putative topologically associated domain (TAD) indicating potential interaction between the *GHSR* locus and the candidate CpG-SNP.

## Discussion

Following domestication, long-term artificial selection contributed to the formation of many pig breeds with spectacular phenotypic diversity. In the past few decades, modern breeding approaches imposed intense selective pressures on pig breeds like Duroc, Landrace and Large White for the improvement of desirable traits. These commercial lean-type breeds were primarily sculpted for superior carcass yield and fast growth rate, while Eastern pig breeds (mainly Chinese) were renowned for excellent meat quality, high-fat content, and early maturity. However, the genetic architecture behind phenotypic variation, especially the effects on morphogenesis, was not yet fully understood. Our genome-wide screen for selective sweeps based on seven high-quality genomes and 1,081 individuals, the most extensive dataset so far, enabled exploiting the contributions of coding and regulatory signatures on several important traits, particularly skeletal muscle development. The current work not only establishes a rich resource to probe functional variants based on genomic sequences, gene expression, and DNA methylation data but also could advance our understanding of the mechanistic basis how artificial selection determines phenotypic changes by reshaping the pig genome, transcriptome and epi-genome.

A growing number of studies have explored genome-wide selective sweep signatures during domestication and breeding in pigs^2,3,7-10,32^, but the number of breeds and sample size in these studies remained limited. With the seven assembled genomes and 1,081 individuals, including 78 diverse domestic breeds presented here, we furnished a comprehensive pan-genome and genetic variation resources, especially structural variations representing the abundant genetic diversity in extant breeds, and significantly expanded the catalogue of current genetic information. Our work provides further support for phenotypic and genetic divergence between Eastern and Western pig populations. Intensive breeding practice in several Western breeds consolidates more concentrated regions, whereas selective sweeps in the Eastern pigs showed more dispersive patterns across the genome, which might endured less selective pressure^2^. Besides, EAD and WED pigs only shared a very small number of swept regions and genes, supporting the relatively independent process of domestication and breeding of these two categories^1^. The rich repertoire of promising candidates provides a valuable resource for the research community and future genome-designed breeding in pigs.

The enrichment/depletion analysis for all putative genetic variations exhibited a significant depletion within exons, indicating strong signals of purifying selection on coding polymorphisms, especially missense mutations with potentially deleterious effects. However, many coding loci were consciously kept and further dispersed in the populations by modern breeding procedures, benefiting from their favourable consequences on several traits of economic interests, like causative mutations in the *MC1R*, *NR6A1*, *PRKAG3*, *RYR1*, and *SERPINA6* genes^7,19,22,33,34^. In addition to these well-known genes, we identified two promising candidate genes, *BRCA1* and *ABCA3*, by bioinformatic and experimental approaches. The *BRCA1* gene codes for a tumor suppressor whose mutations in humans were associated with a predisposition to breast and ovarian cancers due to abnormal cell cycle pathway^35,36^. We observed remarkably strong signals of selective sweeps in the *BRCA1* gene in the WED group. Interestingly, *BRCA1* was also shown to be under selection during chicken domestication, especially in commercial broiler breeding^37^. This consistent finding reflected the convergent patterns in the human-driven selection process on specific preferred breeding goals for farm animals, like higher meat production and faster muscle growth. Porcine enzootic pneumonia (PEP) was a chronic and highly contagious respiratory disease that causes considerable economic losses and is primarily engendered by the Mhp^25,26^. Generally, Chinese local breeds showed a higher susceptibility to Mhp than Western lean-type breeds. Our results indicated that the mutant T allele results in a non-synonymous substitution leading to reduced alveolar macrophage and further attenuated host defense, which was supported by *in vitro* Mhp infection experiments in cell culture. Besides, we also identified several swept genes with coding mutations, like *ALS2*, *EXOC2*, *PRDM15*, and *SEMA3B/E*, which played an essential role in muscle growth, fat deposition, and neuronal development^8,38-40^. Altogether, these functional coding variants constitute valuable information for future pig breeding.

Divergent artificial selection has resulted in a tremendous difference in meat production between Chinese native and Western lean-type pig breeds, but the developmental programs underlying this discrepancy were unclear. We reasoned that the phenotypic consequence was attributed to the timing of the myogenesis process. Heterochrony was broadly defined as the genetically controlled changes in the rate or timing of developmental events in an organism compared to its ancestors or other organisms, which could lead to a tremendous difference in size, shape, characteristics of certain organs and features^13,28,41^. A variety of studies have explored the molecular heterochrony, mainly in the brain and retina across several species and identified numerous hub genes driving organic development, morphogenesis, and evolution^42-46^. It has been widely accepted that muscle growth can be divided into two major periods, prenatal myogenesis determined by an increase in myofibres number (hyperplasia), and postnatal growth, which was achieved by augmenting the size (hypertrophy) of myofibres^47^. Despite the high similarity, our results demonstrated that LDR show a relatively more prolonged timing of myofiber formation in the prenatal stages than TC, but a faster muscle growth and greater lean accretion in the postnatal period. We found more differentially progressing genes and greater overlap with putative swept candidates in the prenatal stages, implying that the effects of artificial selection on skeletal muscle development can be traced back to the very early stages of myogenesis. Together, our data suggested that the vast difference in pork production between fat- and lean-type pig breeds was attributed to the heterochrony in skeletal muscle development, and artificial selection had served as the crucial driving force in muscle growth, especially for prenatal hyperplasia.

DNA methylation was an epigenetic mark frequently occurring at CpG dinucleotides which plays an important role in shaping phenotypic diversity in the domestication of animals^48,49^. In line with previous findings^48-51^, we found that Landrace pigs displayed more hypermethylated regions than TC pigs, resulting from extensive breeding. It appears that selective sweeps are more likely to overlap with DMRs detected at multiple stages, implying that surrounding fixed alleles might have a persistent effect on DNA methylation variation. Additionally, heterochronic genes, especially those covered by selective sweeps, were prone to being modulated by DMRs, suggesting sites with differential methylation levels were vital for controlling developmental progression. This finding provided novel insight into the spatial-temporal patterns behind how artificial selection sculpts organogenesis by reshaping the epigenome.

In general, methylated cytosines exhibited a higher mutation rate, and many studies have reported a significant association between CpG to TpG/CpA transitions and developmental disorders^52,53^. The majority of genetic underpinnings under selection were located within non-coding regions and generally act as regulatory elements to govern gene expression^54-56^. We identified an insertion as a potential enhancer regulating *BDH1* in Eastern pigs, which resulted in the difference in myofiber proliferation and differentiation between Eastern and Western breeds. BDH1 was an important rate-limiting enzyme responsible for ketone metabolism and ATP synthesis^57,58^. The higher expression of *BDH1* induced by the insertion might play a key role in myofiber proliferation and differentiation. Besides structural variations, the present studyed shows that (nearly) fixed SNPs were important sources for inter-breed differences in methylation patterns and can serve as regulatory sequence variants correlated with traits of interest. As a strong candidate under selection, *GHSR* was commonly associated with growth and carcass traits in livestock^59,60^, which was also supported by our association results of the regulatory SNP (chr13_111051076_C/T). Furthermore, higher GHSR and Ghrelin concentrations during pregnancy would lead to higher fetal weight and postnatal weight gain^61-63^. Interestingly, we found that the *GHSR* gene displayed a more advanced expression pattern only in the prenatal stage in LDR pigs, which may be the result of altered DNA methylation status. This finding means that administration of ghrelin happens at the earlier pregnancy and lasts longer in LDR, hence resulting in greater muscle mass and body weight. These data demonstrate the importance of pigs as a biomedical model to probe molecular mechanisms underlying some human metabolic disorders like obesity and diabetes.

## Methods

### Sample collection and sequencing

To capture the full genetic diversity of pigs around the world, we collected the ear tissues of 313 individuals from 30 distinct breeds, representing populations at the climatic and geographical extremes of China. All experiments with pigs were conducted under the guidance of ethical regulation from the Chinese Academy of Agricultural Sciences, China. Sequencing data for a total of 98 individuals from 11 native breeds were shared by our collaborators (unpublished data). In addition, we downloaded 670 high-depth whole-genome sequencing data from the Sequence Read Archive (SRA) database (Supplementary Table 3) to establish the most comprehensive pig genome sequence dataset so far. Genomic DNA was extracted from the ear tissues of 313 samples using a standard phenol-chloroform method. About 1.5 μg DNA was used to construct an approximately 350 bp insert size DNA library at Novogene (Tianjin, China). In brief, the DNA sample was fragmented, then the ends were repaired and ligated to the adaptor. Next, adapter-ligated DNA was selected by running a 2% agarose gel to recover the target fragments. Polymerase chain reaction (PCR) amplification and purification were then performed. According to the standard manufacturer’s instructions, the quantified library was sequenced on the Illumina NovaSeq platform (Illumina, CA, USA). The raw sequences were cleaned to remove adaptors and sequencing errors. Reads containing the sequencing adaptor, more than 10% unknown nucleotides, or more than 50% bases of low quality were removed (quality scores in the Phred scale less than 5). The skeletal muscle (*longissimus dorsi*) samples were newly collected from Tongcheng pigs at 27 developmental stages, including embryonic days 33, 40, 45, 50, 55, 60, 65, 70, 75, 80, 85, 90, 95, 100 and 105 (abbreviated as E33, E40, E45, E50, E55, E60, E65, E70, E75, E80, E85, E90, E95, E100 and E105) and postnatal days 0, 9, 20, 30, 40, 60, 80, 100, 120, 140, 160 and 180 (abbreviated as D0, D9, D20, D30, D40, D60, D80, D100, D120, D140, D160 and D180). Total RNA for RNA-seq and DNA for WGBS was extracted using the standard procedures and were sequenced on the Illumina HiSeq X Ten platform (Illumina, CA, USA). Both WGBS and RNA-seq libraries were performed in three biological replicates at each developmental stage. The time-course transcriptome and methylome data from Landrace pigs were obtained from our previous study^27^.

### Genome assembly

The genomic DNA libraries from seven individuals were constructed and sequenced using the PacBio Sequel platform with CCS mode. Hifiasm^64^ was used to generate the assembly from HiFi CCS reads using default parameters. Hi-C fragment libraries from the same samples were generated with insert sizes of 300-700 bp and sequenced on the Illumina platform. The enzyme used in the Hi-C library was DpnII which cuts DNA at “GATC”. Adapter sequences of raw reads were trimmed, and low-quality paired-end reads were removed for clean reads by using the fastp^65^ program. Bowtie2^66^ was used to align the clean reads to the assembled contigs. We filtered low-quality reads using a HiC-Pro pipeline^67^ with the default parameters. The valid reads were used to anchor chromosomes with Juicer^68^ and 3D-DNA pipeline^69^. The assembly completeness was assessed using the Benchmarking Universal Single-Copy Orthologs (BUSCO) program^70^ containing the Mammalia odb10 gene set (9,226 BUSCO genes). The single-copy plus duplicated complete BUSCO gene counts were reported.

### Pan-genome construction

Seven HiFi assemblies and our previous Luchuan genome were aligned to the Duroc pig reference genome (Sscrofa11.1) by employing minimap2 (-a -x asm10)^71^. A reliable alignment was defined as a continuous alignment longer than 300 bp with sequence identity higher than 90%. Sequences with no reliable alignments were kept as unaligned sequences. Next, MUMmer^72^ was used to map unaligned sequences to the Sscrofa11.1 genome using the parameters -maxmatch and then used the delta-filter -I 90 -l 300 -1 parameter with the one-to-one alignment block option to filter the alignment results. The resulting sequences were aligned against the Sscrofa11.1 genome using blastn (with the parameters-word_size 20-max_hsps 1-max_target_seqs 1-dust no-soft_masking false-evalue 0.00001)^73^, and the sequences showing identity >90% to Sscrofa11.1 sequences were removed. The remaining sequences were merged according to the adjacent regions within 200 bp, and sequences of < 500 bp in length were removed. Next, the generated sequences were aligned to the GenBank nt database with BLASTN^73^ with parameters ‘-evalue 1e-5-best_hit_overhang 0.25 -perc_identity 0.5 -max_target_seqs 1’. Sequences with best hits from other species, or covered by known animal mitochondrial genomes, were possible contaminations and removed. Subsequently, the remaining sequences obtained from all the assemblies were combined. To remove redundancies, all-versus-all alignments with minimap2^71^ (-X -a -x asm10) were carried out.

### Annotation of transposable elements

The transposable elements were identified using two methods: *de novo* repeat identification and known repeat searching against existing databases. RepeatModeler (http://www.repeatmasker.org/RepeatModeler/) was used to predict repeat sequences in the genome, RepeatMasker^74^ was then used to search the genome against the *de novo* transposable element (TE) library. RepeatMasker^74^ and the Repbase^75^ database were used to identify known transposable element TE repeats in the assembled genome. RepeatMasker was applied for DNA-level identification and RepeatProteinMasker was used to perform protein-level identification.

### Gene prediction

For the homolog prediction method, the genome sequences and annotation files from six mammalian genomes, including human, mouse, cattle, dog, goat, and Duroc pig, were downloaded from Ensembl database (Ensembl release 105). Besides, the Luchuan pig protein sequences were downloaded from the China National GenBank (CNGB; https://db.cngb.org/) under the accession of CNP0001159. These sequences were aligned to the genome assembly using GeMoMa^76^ to detect homologous peptides. For the RNA-seq-based prediction approach, the raw Illumina short reads of nine tissues were used as well as one pool RNA library (i.e. subcutaneous adipose, kidney, heart, lung, longissimus dorsi muscle, liver, psoas major muscle, spleen, ovary. Accession numbers: SRR3160015, SRR3160012, SRR3160008, SRR3160011, SRR3160014, SRR3160009, SRR3160017, SRR3160013, SRR3160010, SRR3160016) were downloaded from NCBI for further analyses. All raw reads were assessed using fastp^65^ with the default setting. The clean reads were aligned to the genome assembly using HISAT2^77^ to identify putative exon regions and splice junctions. StringTie^78^ was then used to assemble the mapped reads into gene models and validated by Program to Assemble Spliced Alignment (PASA)^79^. Genes that had PASA support with correct structure but were lost in homology-based prediction were added into the gene set. Finally, untranslated regions and alternative splicing regions were determined using PASA^80^.

### Gene functional annotation

Five public databases, including NCBI non-redundant protein sequence database, SwissProt^81^, Kyoto Encyclopedia of Genes and Genomes (KEGG)^82^, Translation of European Molecular Biology Laboratory and Gene Ontology (GO), were used for functional annotation of the reference gene set. Putative domains and GO terms of these genes were identified using InterProScan^83^, while the Diamond program^84^ was used to compare the protein sequences of the pig genome against the remaining four public databases with an E-value cutoff of 1e-05.

### SV detection

The minimap2^71^ tool with the parameter “-x splice -t 10 -k 12 -a -p 0.4 -N 20” was used to align the eight individual CDS to the pig reference genome Sscrofa11.1. Subsequently, anchorwave^85^ was performed with the setting “genoAli-IV”. The output maf files were converted to sam format with maf-convert.py (https://gitlab.com/mcfrith/last/-/blob/main/bin/maf-convert). Finally, samtools mpileup^86^ was employed to call SVs.

### Reads mapping of whole-genome sequencing data to the pan-genome

We merged putative non-reference pan-sequences and the Sscrofa11.1 reference assembly as the pig pan-genome. The fastq reads were first trimmed by fastp^87^ with default parameters. Next, all clean reads, including our newly generated samples, were aligned to the pan-genome using the BWA-MEM pipeline. The mapped reads were then sorted, and duplicates were removed by Picard tools (https://broadinstitute.github.io/picard/) and SAMtools^88^.

### Genome-wide screening of SNPs and INDELs

The genome-wide variants were called for each sample by the GATK UnifiedGenotyper^89^ with *- glm BOTH -rf BadCigar --sample_ploidy 2* option. To ensure high accuracy of variants calling, SNPs with *QD* < *2.0 || FS* > *60.0 || MQ* < *20.0 || MQRankSum* < *-12.5 || ReadPosRankSum* < *- 8.0* were filtered. Gene-based SNP annotation was performed according to the annotation of the pan-genome using the package ANNOVAR^90^ and Ensembl Variant Effect Predictor^91^. Based on the genome annotation, SNPs were categorized as occurring in exonic regions, 5’ or 3’ untranslated regions, intronic regions, splicing sites (within 2 bp of a splicing junction), upstream and downstream regions (within a 1 kb region upstream or downstream from the transcription start site), or intergenic regions. SNPs in coding exons were further grouped as either synonymous SNPs or nonsynonymous SNPs. To assess the sequence conservation of missense mutations, we first downloaded the amino acid sequences of nine related mammals from the NCBI or Ensembl database, including human, mouse, cattle, horse, goat, tiger, cat, dog, and rabbit. Subsequently, multiple sequence alignments were performed by the MUSCLE program^92^. Meanwhile, the SWISS-MODEL workspace^93^ was used to predict protein structure models for wild-type and mutant variants.

### Phylogenetic and population genetic analyses

To provide insight into phylogenetic relationships among different pig breeds, we performed a comprehensive genomic survey based on all autosomal high-quality bi-allelic SNPs with a call rate ≥ 90% and a minor allele frequency ≥ 5%. A neighbor-joining tree was constructed using the program TreeBeST (http://treesoft.sourceforge.net/treebest.shtml) with 200 bootstrap replicates. The tree was displayed using Interactive Tree Of Life (iTOL)^94^. To infer the population structure, we used ADMIXTURE^95^, which implements a block-relaxation algorithm. We also filtered SNPs by testing Hardy-Weinberg equilibrium (HWE) violations (*p*-value > 10^-4^) and reconstructed the model-based clustering analysis. To identify the best genetic clusters K, cross-validation error was tested for each K value from 2 to 10. The principal component analysis (PCA) was conducted using the program GTAC^96^. The pattern of linkage disequilibrium (LD) for these regions of interests was computed by using the LDBlockShow software^97^.

### Selective sweeps analysis

We first excluded the potential mixed samples between the Eastern and the Western pigs according to the results of population structure, to ensure the accuracy of the putative selective sweeps. When K = 2, if the lineage of a Western pig is more than 20% for an Eastern breed, it will be filtered out, and vice versa for Western breeds. SNPs with minor allele frequency below 5% were removed from this analysis. Subsequently, a computationally advanced composite-likelihood ratio (CLR) test was used to detect genome-wide selective sweeps by SweeD software^98^. CLR scores were computed for each 10-kb non-overlapping window along all the autosomes. Finally, the top 1% of windows (with the highest CLR scores) were considered to be candidate selective regions. To evaluate the association between candidate variants and traits of economic importance in pigs, we leveraged genotype and phenotype data of 589 F2 individuals from our collaborator, which was derived from a cross between five Large White (European, WED) boars and sixteen Min (Chinese, EAD) sows^23^. We first sorted the pedigree information according to the birth order by a custom python script, and next established the relative kinship coefficients matrix. Finally, we compared the differences in the slaughter weight at 240 days and lean meat proportion using a mixed linear model (MLM) with the matrix as the covariate. The *p*-value was corrected by Bonferroni approach, and a corrected *p*-value of less than 0.05 was regarded as significantly different. To classify functional categories of putative selective sweeps in detail, we estimated the Jaccard index between these regions and five genomic features with 1,000 permutations. GO enrichment analysis of swept genes was implemented with the Metascape tool^99^. GO terms with corrected *p*-values < 0.05 were considered significantly enriched. To explore enriched phenotypes driven by putative selection signatures, we performed trait/QTL enrichment analysis by hypergeometric test against the pig QTL database^21^. We focused on the QTLs with confidence interval less than 1 Mb, given that the QTL confidence intervals are too large to be used efficiently in the post-processing.

### Whole transcriptome sequencing

Total RNA was extracted with TRIzol Reagent (Invitrogen, Shanghai, China). A total amount of 3 μg RNA per sample was used as input material for the RNA sample preparations. Ribosomal RNA was removed by Epicentre Ribo-zeroTM rRNA Removal Kit (Epicentre, USA), and rRNA-free residue was cleaned up by ethanol precipitation. The RNA concentration was monitored with a Qubit Fluorometer (Invitrogen), and the RNA quality was evaluated by the Agilent Bioanalyzer 2100 system (Agilent Technologies, CA, USA) prior to library preparation. Subsequently, sequencing libraries were generated using the rRNA-depleted RNA by NEBNext UltraTM Directional RNA Library Prep Kit for Illumina (NEB, USA) following the manufacturer’s recommendations. Finally, paired-end sequencing with 150 nucleotides at each end was implemented on the Illumina NovaSeq 6000 platform (Illumina, CA, USA).

### Whole-genome bisulfite sequencing

A total amount of 100 ng genomic DNA spiked with 0.5 ng lambda DNA was fragmented by sonication to 200-300 bp with Covaris S220. These DNA fragments were treated with bisulfite using EZ DNA Methylation-GoldTM Kit (Zymo Research), and the library was constructed by Novogene Corporation (Beijing, China). Subsequently, library quality was assessed on the Agilent Bioanalyzer 2100 system, and 150 bp pair-end sequencing of each sample was performed on the Illumina Novaseq 6000 platform (Illumina, CA, USA).

### RNA-seq analysis

The raw data was filtered by fastp with default parameters^87^. The clean data were mapped to the constructed pan-genome and annotation files with STAR aligner^100^. Gene-level read counts were enumerated at the same time, and used as input for DESeq2 package^101^. Based on these normalized counts by rlogTransformation function in DESeq2 software, we conducted PCA, hierarchical clustering and k-means clustering. Subsequently, we applied TimeMeter tool^28^ to evaluate temporal gene expression similarity and detect differentially progressing genes between Landrace (LDR) and Tongcheng (TC) breeds. Positive progression advance scores (PAS) indicated faster expression patterns in TC (TC acceleration genes), while negative means accelerated changes in LDR (LDR acceleration genes). GO term of Biological Process analysis predicted with DEGs was conducted with the Metascape^99^.

### Whole-genome bisulfite sequencing analysis

Our built pan-genome was firstly transformed to Bisulfite Genome with Bismark tool (https://www.bioinformatics.babraham.ac.uk/projects/bismark/). Then the clean data after quality control were aligned to the Bisulfite Genome with Bismark based on the default parameters. The methylation information for each cytosine site was extracted after filtering the duplicate reads. Differentially methylated cytosine sites (DMCs) and regions (DMRs) were identified using MethylKit software^102^. For the DMCs analysis, the cytosine sites with coverage less than ten were removed. A sliding window approach was used to calculate DMRs, in which both the window and the step were set to 1,000 bp, cytosine sites with coverage less than five were removed and regions that contained at least three cytosine sites were left for the downstream analyses. Both the DMCs and DMRs were defined with the criterion of Bonferroni correction *q*-value < 0.05 and meth.diff > 30. We focused on the gene body region (from TSS to TES), promoter region (upstream and downstream 2 kb from the TSS) and transposon element to assess the methylated information annotation.

### Correlation analysis between gene expression and DNA methylation

Before correlation analysis, we removed genes with rlog-normalized counts less than zero and more than 20. Then, the DNA methylation levels in the gene body and promoter region for each expressed gene were computed. The Pearson correlation coefficients (r) between DNA methylation levels and gene expression of different features were calculated in R software.

### Isolation of porcine adipose cells

The subcutaneous adipose tissues of the neck and back from newly born pigs were collected under the aseptic state and washed with PBS buffer containing a high concentration of penicillin and streptomycin three times. The visible blood vessels and connective tissues were cut off. Subsequently, the adipose tissues were cut into small tissue blocks of 1 mm^3^, and type I collagenase digestion solution was added for digestion for 1 h (37℃, vibrating every 5 min). Then the digestible was filtered with a double-layer nylon sieve of 100 μm and 25 μm, and the filtrate was centrifuged at 1500 r/min for 10 min. After centrifugation at 1,000 r/min for 5 min, the supernatant was discarded, and the serum-free culture medium was added. The culture medium and the cell mixture were mixed evenly, centrifuged at 1,000 r/min for 5 min, then the supernatant was discarded, and the complete medium was added.

### Cell culture

Mouse C2C12 myoblasts and HEK-293T cells were obtained from Peking Union Medical College Hospital and cultured in Dulbecco’s Modified Eagle’s Medium (DMEM, Gibco, USA) supplemented with 10% fetal bovine serum and 1% penicillin-streptomycin (PS, Thermo Scientific, USA). In addition, porcine adipose cells were cultured in DMEM/F12 -Dulbecco’s Modified Eagle Medium (DMEM/F12, Gibco, USA) supplemented with 10% fetal bovine serum and 1% penicillin-streptomycin (PS, Thermo Scientific, USA). All the cells were cultured at 37°C in a 5% CO2 incubator. The culture medium of C2C12 myoblasts was replaced with DMEM supplemented with 2% horse serum (Gibco, USA) and 1% PS at 80% confluence to induce myogenic differentiation.

### Plasmid construction

The full-length coding sequence regions of the mouse *CD36*, *GHSR* and *NCAPG* genes were amplified by polymerase chain reaction (PCR) using forward and reverse primers containing BamH I and Xho I sites, respectively. The PCR products were inserted into the pcDNA3.1(+) vector (Invitrogen, Shanghai, China). To verify the dual-luciferase activity of candidate SNPs from the *CD36*, *GHSR* and *NCAPG* genes, 400-bp DNA fragments centring on the SNPs were first amplified and then inserted into the PGL4.23 Dual-Luciferase Expression Vector (Promega, Madison, WI, USA). Primers were designed using Primer Premier5 and listed in Supplementary Table 21.

### RNA interference

Based on the CDS region of the target gene provided by NCBI, siRNAs of the target gene were designed using the siDirect website (http://sidirect2.rnai.jp/) and synthesized at GenePharma (Shanghai, China). The siRNA sequences are listed in Supplementary Table 22.

### Cell transfection and Dual-luciferase reporter assay

After 12 h of cell culture, the C2C12 myoblasts or HEK293T cells were transfected with the appropriate plasmids or oligos using Lipofectamine 3000 and Opti-MEM according to the manufacturer’s protocols. These cells were collected after a transfection period of 24 h. A Dual-Luciferase Reporter Assay System (Promega) was used to quantify luciferase activities following the manufacturer’s instructions. Firefly luciferase activity was normalized to Renilla luciferase activity.

### Quantitative RT-PCR (qRT-PCR)

Total RNA was isolated from cultured cells using TRIzol reagent (Invitrogen, USA), and then reverse-transcribed to cDNA using the PrimeScript™ RT Master Mix (Perfect Real Time) (Takara, Beijing, China) according to the manufacturer’s protocols. qRT-PCR was performed on a StepONE Real-Time PCR System (Applied Biosystems) according to the SYBR Premix Ex TaqTM instructions. Each reaction was performed in a 20 μl reaction mixture containing 10 μl of ChamQ SYBR qPCR Master Mix (Vazyme Q311-02, Nanjing, China), 0.5 μl of gene-specific primers, 2 μl of template cDNA and 7 μl of sterile water. All reactions were repeated three times with cDNA from three independent individuals, and the results were analyzed using the 2^-ΔΔCT^ method. Each experiment was repeated three times.

### 5-Ethynyl-2’-Deoxyuridine (EDU) assay

The C2C12 Myoblasts were seeded in 12-well plates. When the cells grew to a density of 50% confluence, they were transfected with overexpression plasmid, siRNA, or miRNA mimics. After transfection for 48 h, myoblasts were exposed to 50 μM EDU (RiboBio, China) for 2 h at 37℃. Subsequently, the cells were fixed in 4% paraformaldehyde for 30 min, neutralized using 2 mg/ml glycine solution, and then permeabilized by adding 0.5% Triton X-100. A solution containing EDU (Apollo Reaction Cocktail; RiboBio, China) was added and the cells were incubated at room temperature for 30 min. The nuclear stain Hoechst 33342 was then added, and incubation was continued for another 30 min. A fluorescence microscope (DMi8, Leica, German) was used to capture three randomly selected fields to visualize the number of EDU-staining cells.

### Immunofluorescence analysis

Cells were cultured and fixed in 6-well plates with paraformaldehyde, treated with 0.5% Triton, blocked with goat serum for 1 hour, and incubated with anti-MyHC (1:500, DSHB MF20, Shanghai, China) antibodies for 2 h. Next, cells were incubated with goat anti-mouse secondary antibodies. Finally, DAPI (1:1000, Invitrogen D3571, Shanghai, China) was added and the cells were observed using the Leica DMI3000 B microscope (Leica).

### AAV-mediated *in vivo* overexpression of *GHSR* and *CD36* in mice

To validate *in vivo* the function of the *GHSR* and *CD36* genes, we first obtained adeno-associated virus 9 (AAV9) serotypes of pcDNA3.1-GHSR, pcDNA3.1-CD36 or pcDNA3.1-Control from HANBI (Shanghai, China). Meanwhile, 7-week-old mice were obtained from the company Huafukang (Beijing, China). The tibialis anterior (TA) muscle of the right leg in each mouse was injected with AAV9 virus (100 µL at titer ≥ 1×10¹³ vg/ml) with pcDNA3.1-GHSR or pcDNA3.1- CD36, and the left TA muscle was injected with pcDNA3.1-Control. The AAV9 vector-treated mice in triplicate were anesthetized with isoflurane and sacrificed by cervical dislocation to collect muscles. For *GHSR*, we collected TA muscles after 10 days of AAV9 injection. To verify accelerated muscle maturation regulated by CD36, we designed a muscle injury and regeneration model. We first injected AAV9 virus overexpressing CD36 or Control 10 days prior to cardiotoxin (CTX) injection, and then collected TA muscles at days 0, 1, 5 and 10 post-CTX injections. After optimal cutting temperature embedding, frozen sections were fixed in 4% paraformaldehyde overnight, and H&E staining was carried out according to the Hematoxylin-Eosin/HE Staining Kit (Solarbio, Beijing, China).

### Mycoplasma hyopneumoniae challenge experiments

The cell density of porcine alveolar macrophages (PAMs) was adjusted to 1 × 10^4^ cells per well in a 24-well plate and cultured in 0.5 ml of RPMI 1640 medium supplemented with 10% FBS and 2% penicillin-streptomycin solution at 37℃. The cells were washed with PBS to remove any unattached cells after 12 h of culture, and then transfected with 0.4 μg DNA using QIAGEN Transfection Reagent (Cat. No. 301005) according to the manufacturer’s instructions. After 24 h transfection, the cells were transfected again. Subsequently, PAMs with wild-type and mutant alleles were treated by Mycoplasma hyopneumoniae. After 6 h incubation, these infected cells were harvested for qRT-PCR. The PAMs and mycoplasma hyopneumoniae materials were from Institute of Veterinary Medicine, Jiangsu Academy of Agricultural Sciences.

## Data availability

Raw sequencing reads generated by this work, including whole-genome sequencing, RNA-seq, and WGBS data, were deposited in the National Center for Biotechnology Information database under the accession number PRJNA754250. In addition, other sequencing data in this study are downloaded from NCBI Gene Expression Omnibus, and all accession numbers are given as Supplementary Table 9. Analysis codes in this work are available from authors upon request.

## Acknowledgements

We thank Prof. Guojie Zhang (Zhejiang University, China) for his valuable comments and suggestions on this project. This work was supported by the National Natural Science Foundation of China (31830090 to Z.T., 32002150 to G.Y., and 32002151 to L.L.), the Agricultural Science and Technology Innovation Program (CAAS-ZDRW202006 to Z.T.), the Basic and Applied Basic Research Foundation of Guangdong Province (2020B1515120053 to G.Y.), the Shenzhen Science and Technology Innovation Commission (JCYJ20190813114401691 to G.Y.), and the Central Government Guiding Funds for Local Science and Technology Development of China (He-Ke ZY220603 to G.Y.).

## Author contributions

Z.T. and G.Y. designed this project and coordinated research activities. G.Y., L.L., J.L., Yalan Y., L.F., Z.P., Y.N., Lei W., M.C., X.F., Xingzheng L., Z.W., H.H. and P.Y. performed bioinformatics analyses. Z.T., Yilong Y., Liyuan W., X.Q., Yun C. and Y.Z., conducted the experiments. Y.T., Yaosheng C., D.M., Lixian W., L.Z., Yonggang L., Yuying L., Xinjian L., Z.Y., D.L., D.Z., Q.Z., and W.W. collected and provided pig materials or sequencing data. G.Y., Z.T., M.G., L.L., Y.L., L.F., G.E.L., J.J. and Y.G. contributed to writing the manuscript. All authors participated in analyzing and interpreting the data.

## Supplementary Figures and Legends

**Supplementary Fig. 1.**
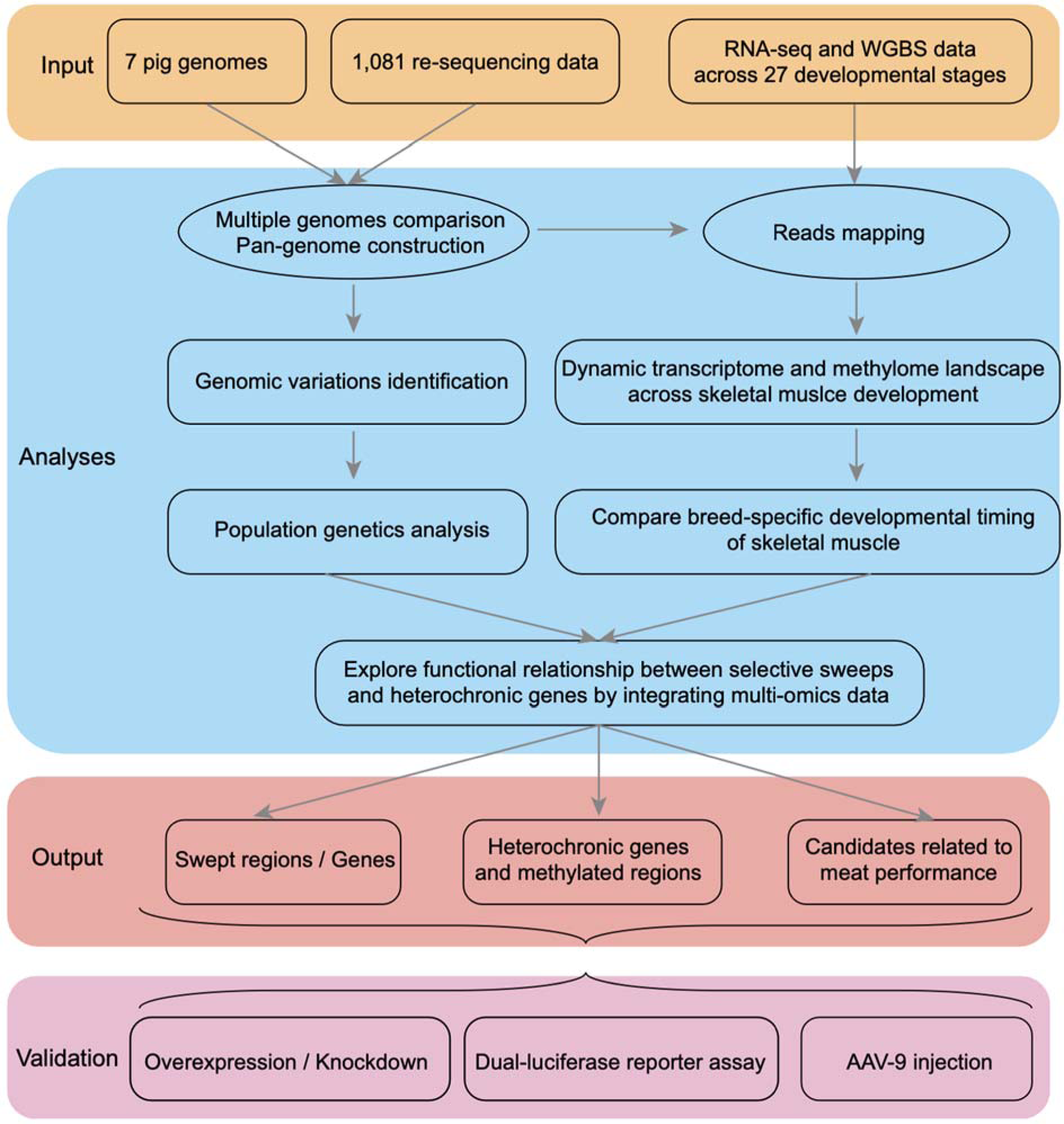
The comprehensive analysis pipeline used in this work.

**Supplementary Fig. 2.**
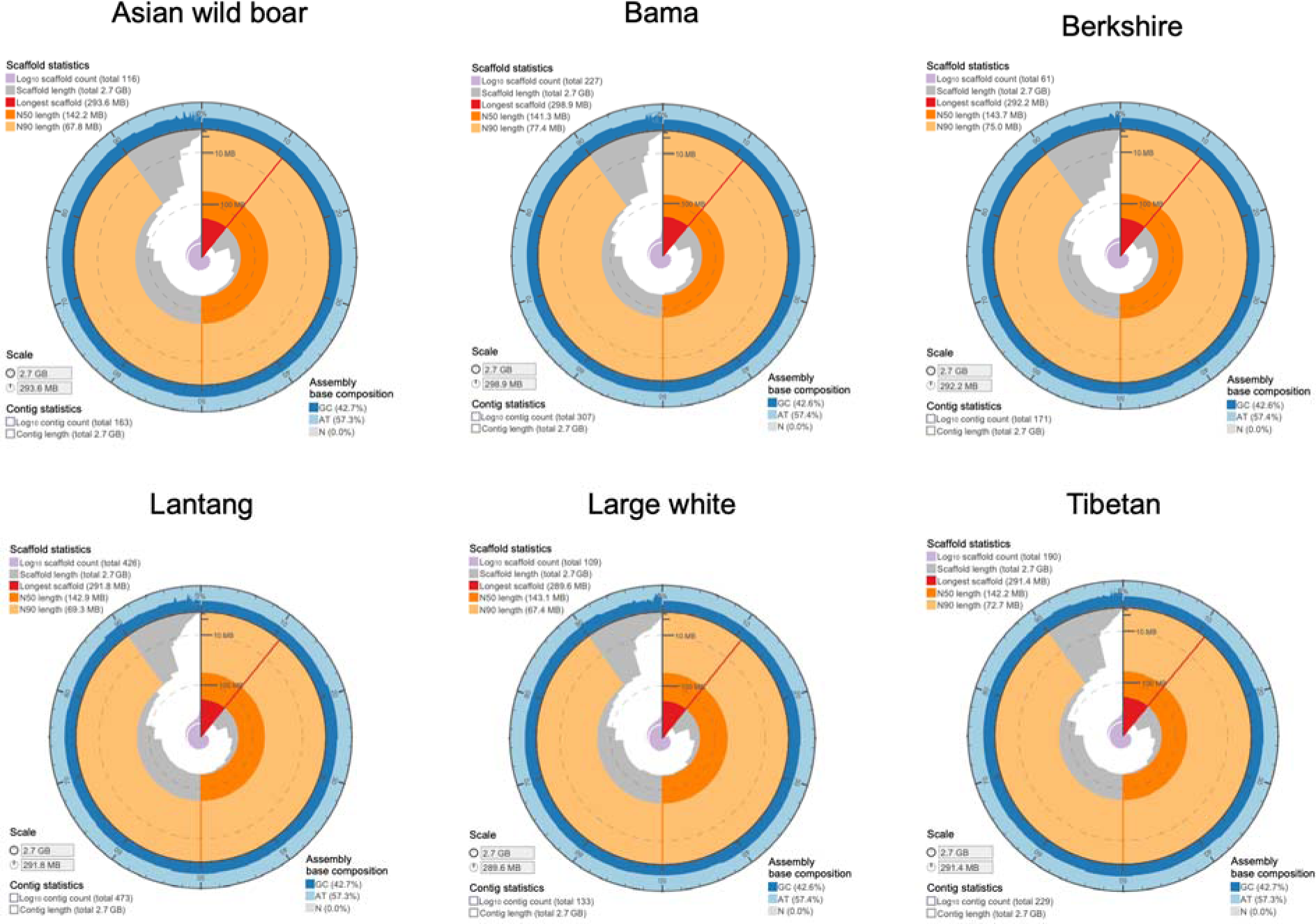
Snail plots showing assembly statistics of six HiFi genomes.

**Supplementary Fig. 3.**
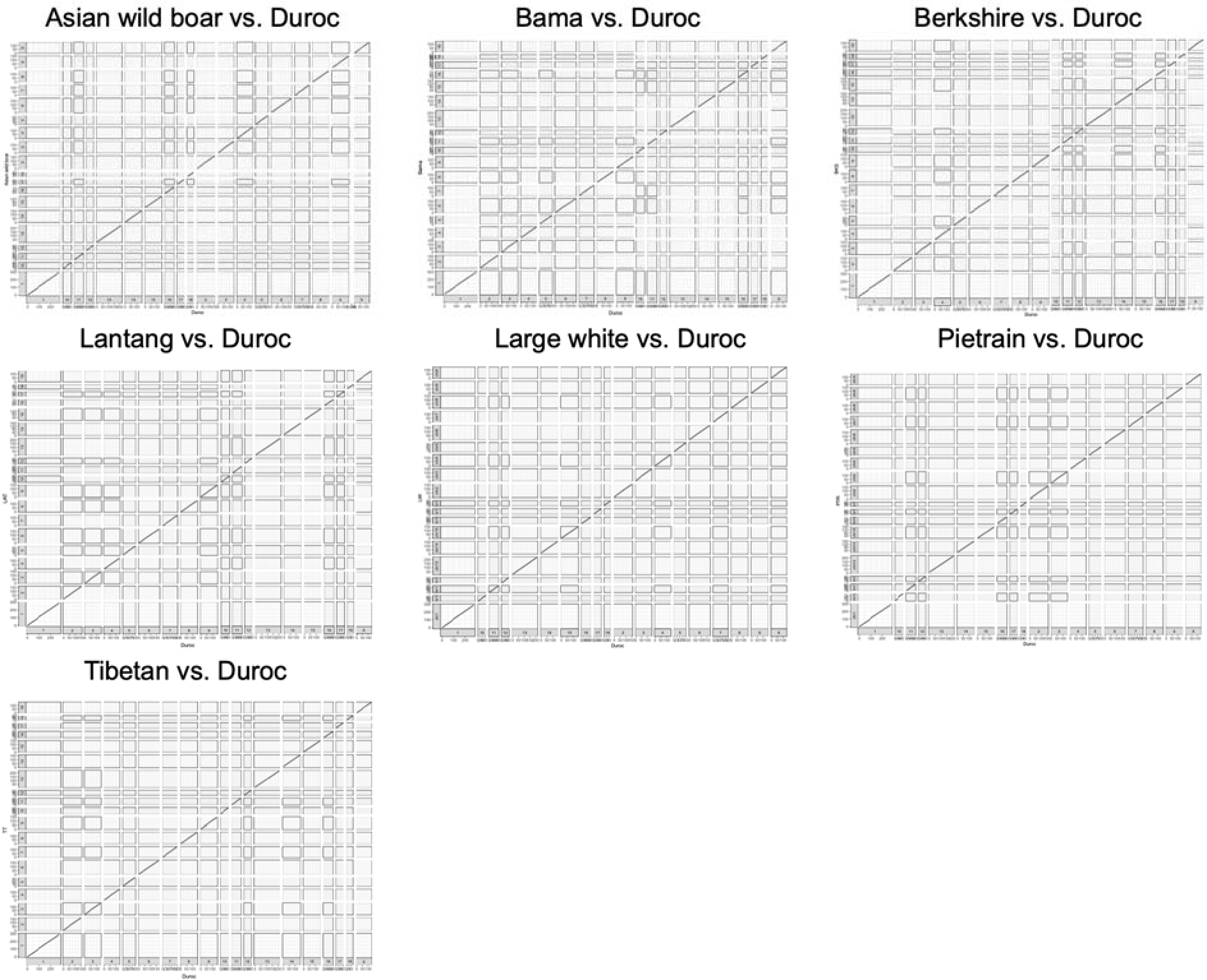
Dotplots showing patterns of synteny and collinearity between assembled genomes and Sscrofa11,1 reference.

**Supplementary Fig. 4.**
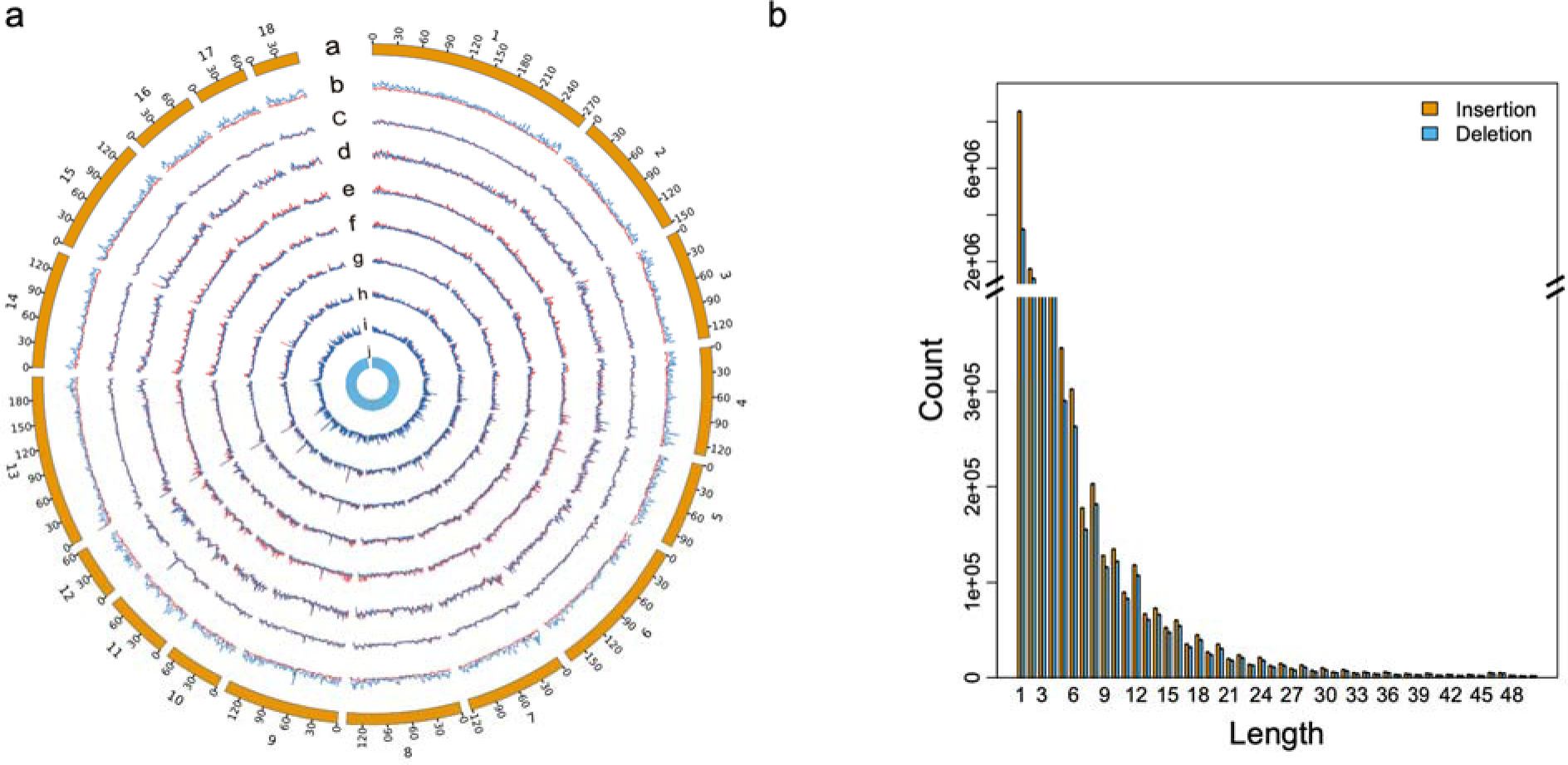
Genome-wide structural variants. **a**, Circos plot of the presence-absence variations including SNPs (red) and SVs (blue) for each pig genome compared with Sscrofa11.1 reference. The innermost blue circle is gene density. The window size is 1Mb. **b**, The length distribution of genomic variations shorter than 50 bp.

**Supplementary Fig. 5.**
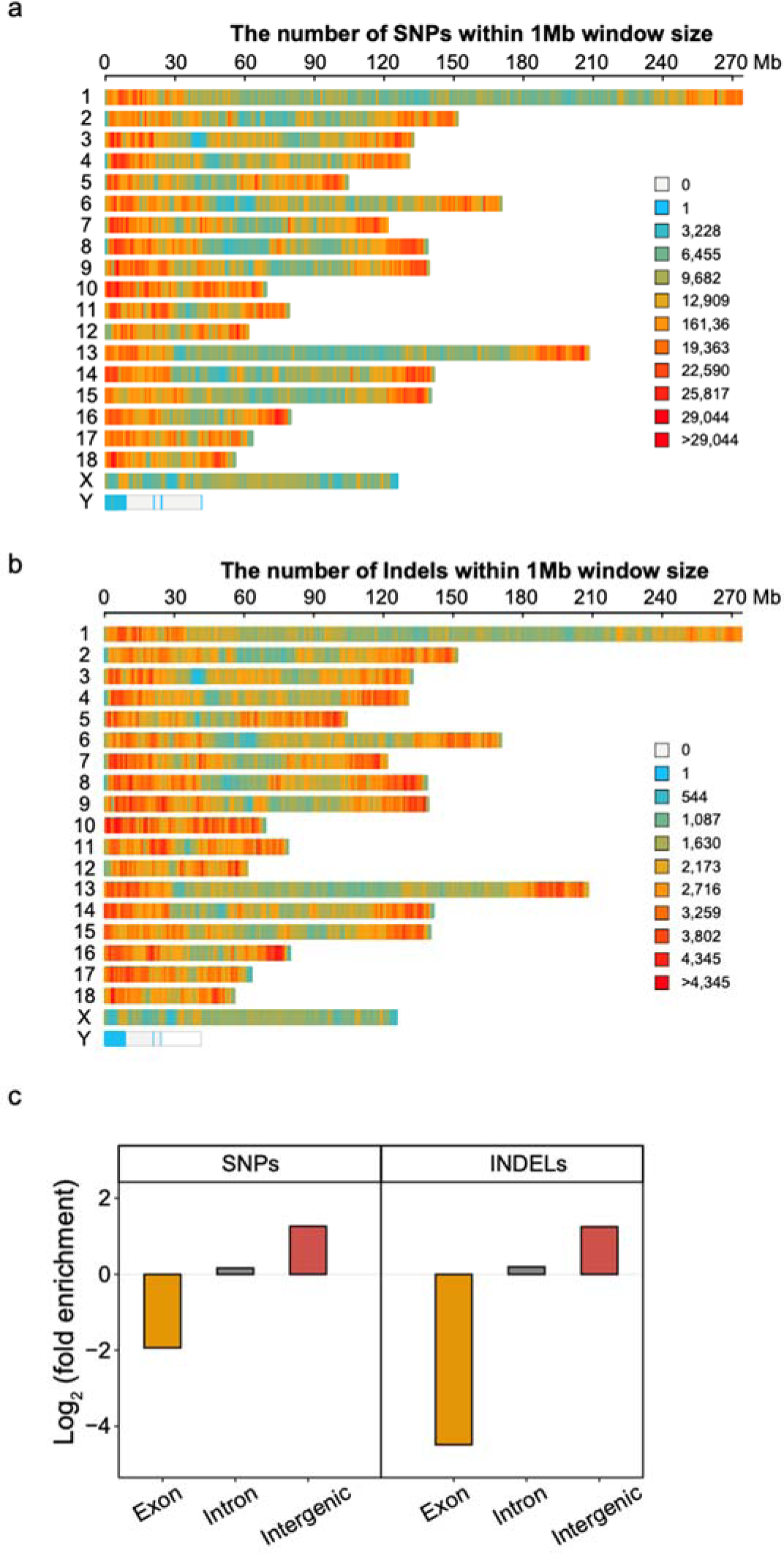
Global map of genetic variations in the pig genome. **a**, Density map of SNPs within 1Mb window size. **b**, Density map of INDELs within 1Mb window size. **c**, Relative enrichment or depletion of SNPs and INDELs within exons, introns and intergenic regions.

**Supplementary Fig. 6.**
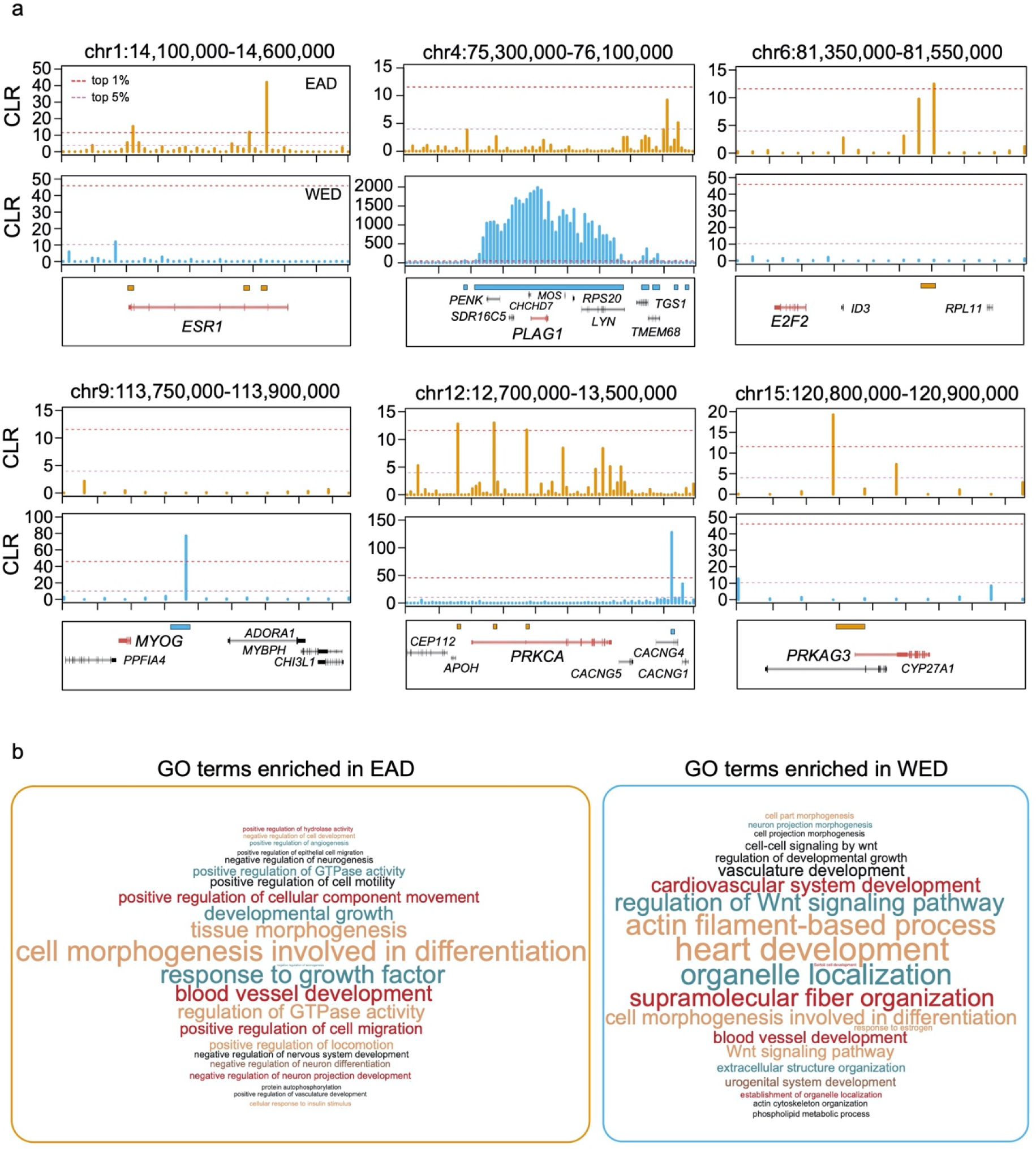
Examples of swept genes in EAD and WED pigs and functional enrichment results. **a**, The composite likelihood ratio (CLR) values of several key genes in EAD and WED pigs. **b**, Word cloud of significant biological process terms (*q*-value < 0.05) based on swept genes in EAD and WED populations.

**Supplementary Fig. 7.**
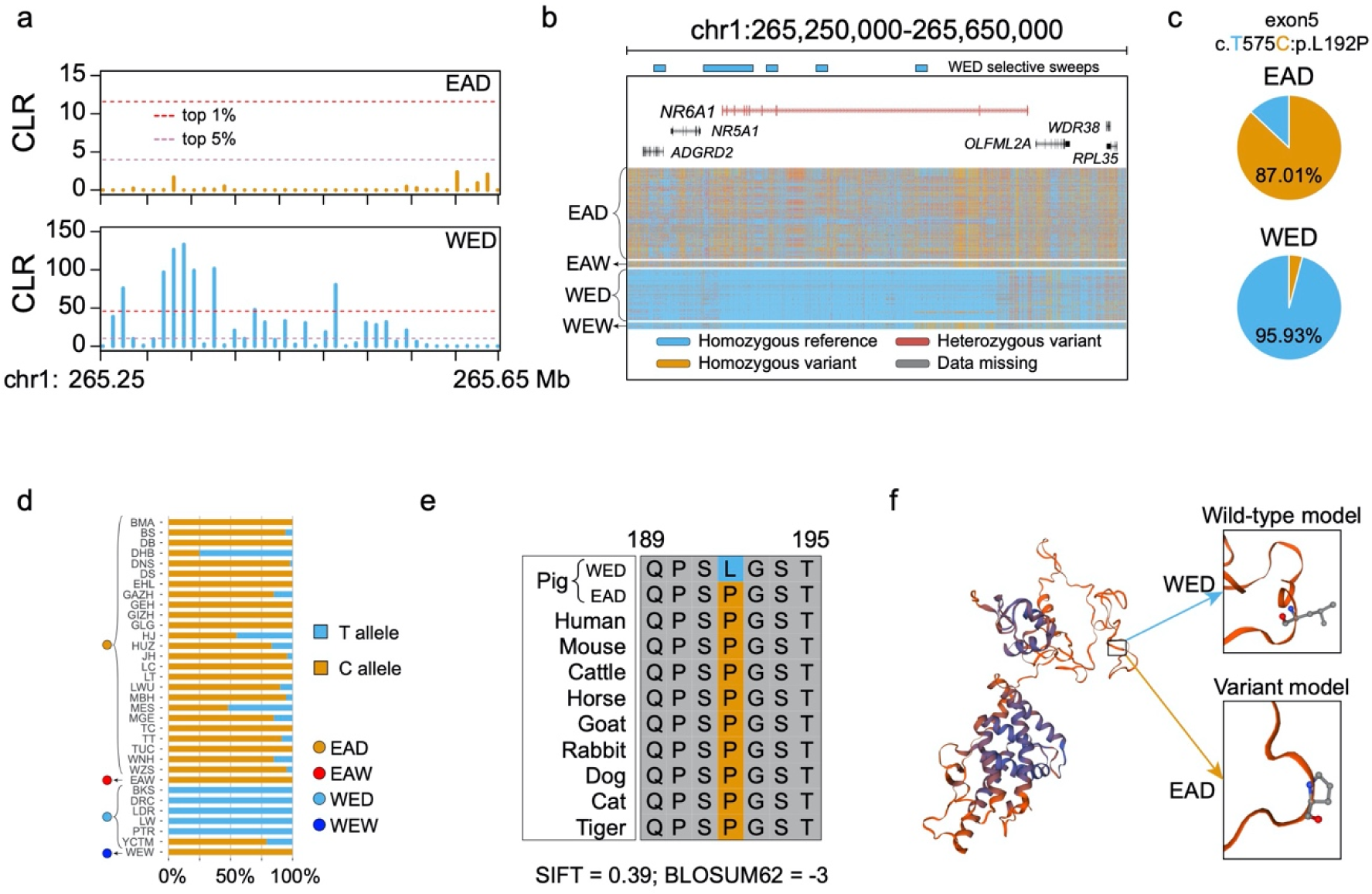
Functional exploration of *NR6A1* gene. **a**, Comparison of CLR values in the *NR6A1* region between EAD and WED pigs. **b**, Genotype spectrum around the *NR6A1* region among different pig populations. **c**, Different allele frequencies of the coding mutation between EAD and WED breeds. **d**, The allele frequency of this variant among different Eurasian pig breeds. BMA, Bama; BS, Baoshan; DB, Debao; DHB, Dahuabai; DNS, Diannan small ear; DS, Dongshan; EHL, Erhualian; GAZH, Guanzhuanghua; GEH, Guangdong litter ear hua; GIZH, Guizhonghua; GLG, Gaoligongshan; HJ, Huanjiang; HUZ, Huai; JH, Jinhua; LC, Luchuan; LT, Lantang; LWU, Laiwu black; MBH, Minbei hua; MES, Meishan; MGE, Mingguang little ear; TC, Tongcheng, TT, Tibetan; TUC, Tunchang; WNH, Wannan black; WZS, Wuzhishan; EAW, Eastern wild boar; BKS; Berkshire; DRC, Duroc; LDR, Landrace; LW, Large white; PTR, Pretriain; YCTM, Yucatan; WEW, Western wild boards. The number of pigs in each breed is greater than 10. **e**, Multi-species alignment of NR6A1 protein for which domestic pigs are fixed or close to fixation for a derived amino acid substitution. This result indicates that Pro is more conserved than Leu. f, Alteration in protein structure resulting from the mutation in the NR6A1 protein. This single amino acid substitution might exert an adverse effect in the protein folding.

**Supplementary Fig. 8.**
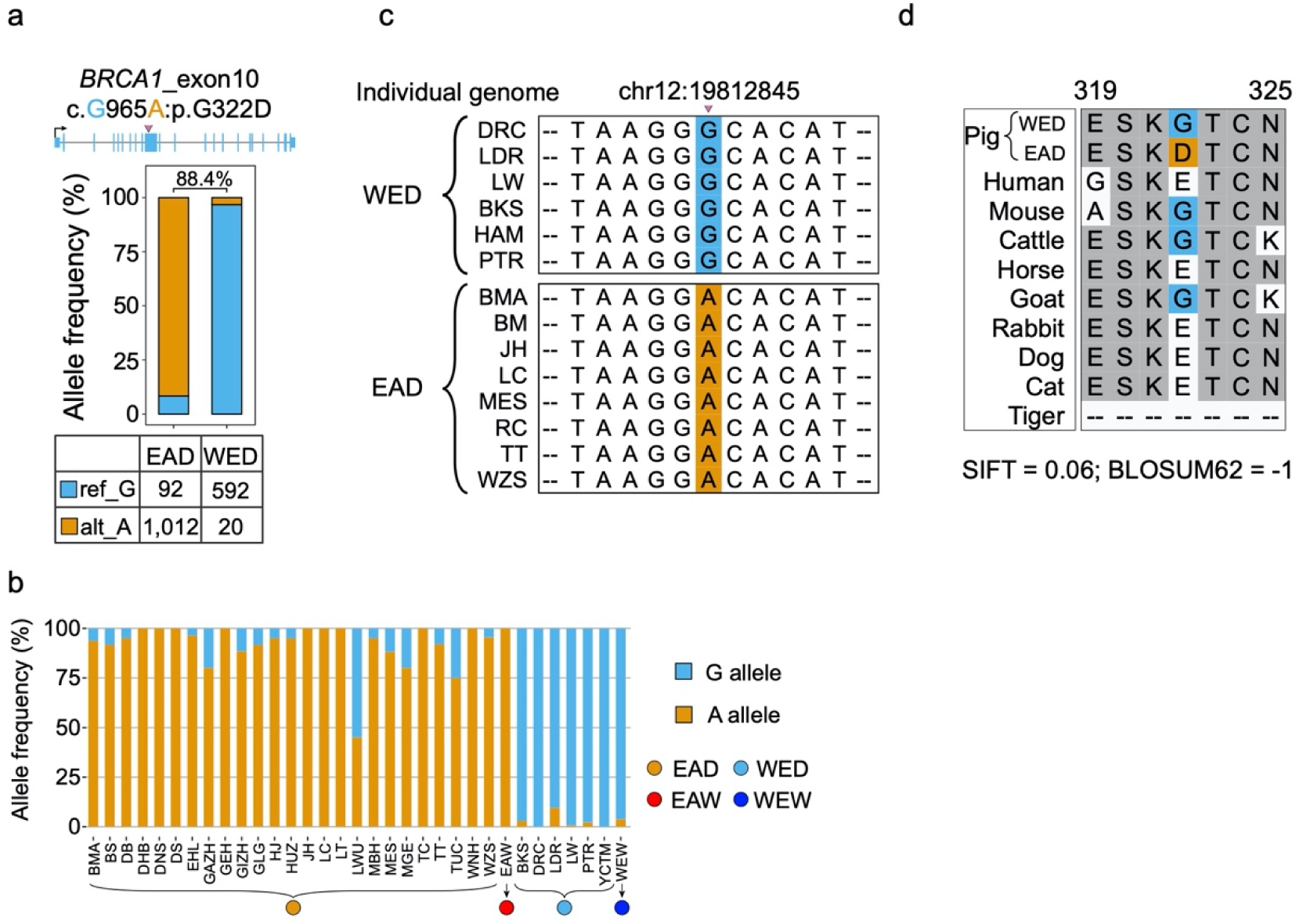
Functional study of the *BRCA1* variant. **a**, The allele frequency of the coding mutation c.G965A in EAD and WED populations. **b**, Comparisons of this variant among different breeds with a sample size greater than 10. The full name for each breed can be found in Supplementary Table 9. **c**, Comparisons of the sequences around this missense mutation across the 14 pig assemblies in pan-genome analysis. **d**, Multiple sequence alignment of amino acids across ten representative mammalian species.

**Supplementary Fig. 9.**
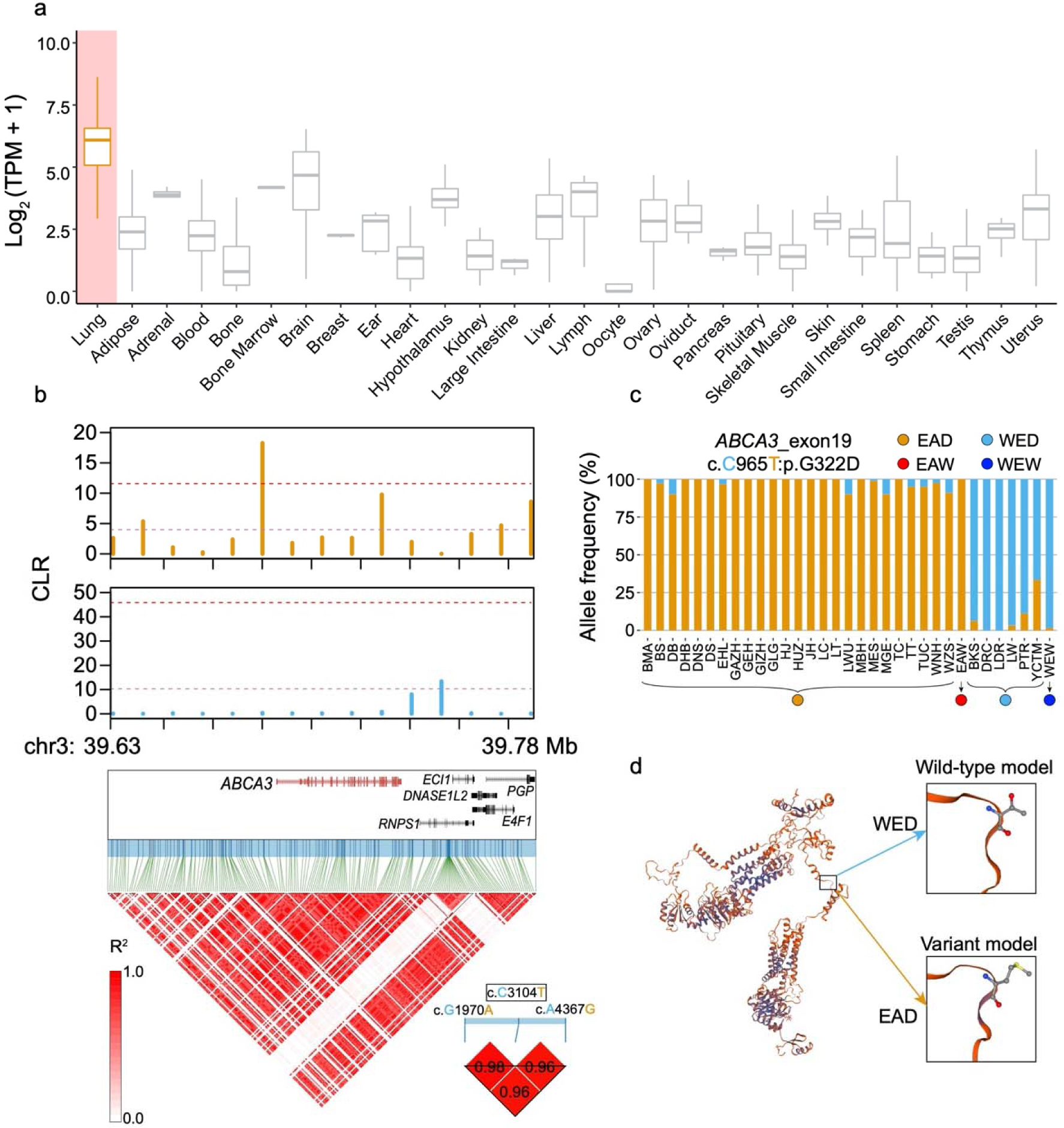
Distinct genomic landscape and functions of the coding variant in the *ABCA3* gene. **a**, The *ABCA3* gene shows predominant expression in lung tissues. **b**, CLR comparison and LD pattern around the *ABCA3* region. **c**, The allele frequency of the mutation c.C965T in *ABCA3* among different pig breeds. The full name for each breed can be found in Supplementary Table 9. **d**, Alteration in protein structure resulting from the mutation in ABCA3 protein.

**Supplementary Fig. 10.**
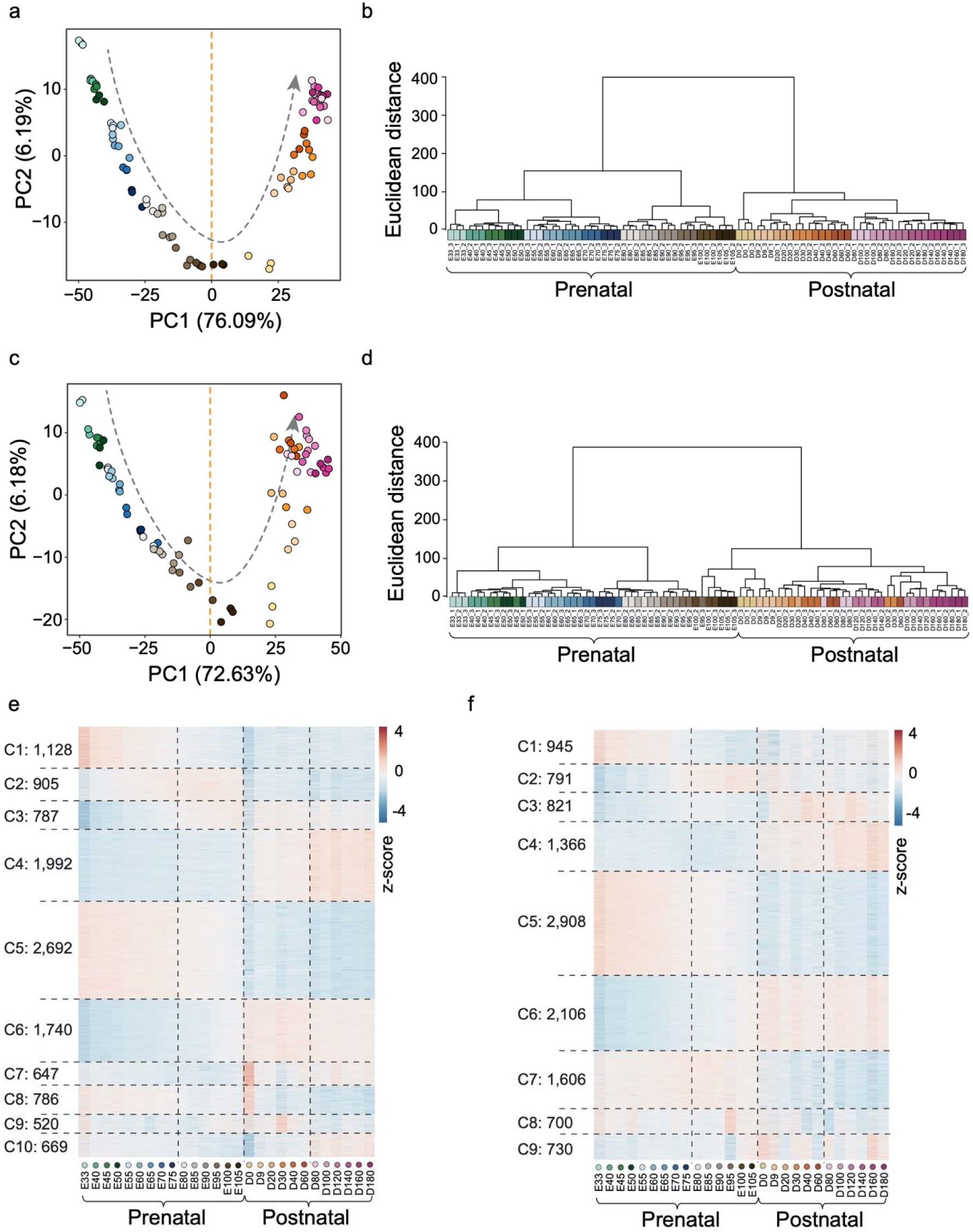
Global transcriptome profiles across skeletal muscle development in Landrace (LDR) and Tongcheng (TC) pigs. **a**, PCA analysis of gene expression data according to top 1,500 variable genes in LDR. **b**, Hierarchical clustering based on the expression levels of top 1,500 variable genes in LDR. Unsupervised PCA and hierarchical clustering results showed the excellent agreement among biological replicates, and a smooth transition from one stage to the neighbouring phase. **c**, PCA analysis of gene expression data according to top 1,500 variable genes in TC. **d**, Hierarchical clustering based on the expression levels of top 1,500 variable genes in TC. **e∼f**, Distinct expression patterns of those dynamic genes by k-means clustering among 27 developmental stages in LDR and TC pigs. By grouping differentially expressed genes (DEGs) with similar transcriptional patterns by k-means clustering approach, we defined nine and 10 clusters in TC and LDR, respectively.

**Supplementary Fig. 11.**
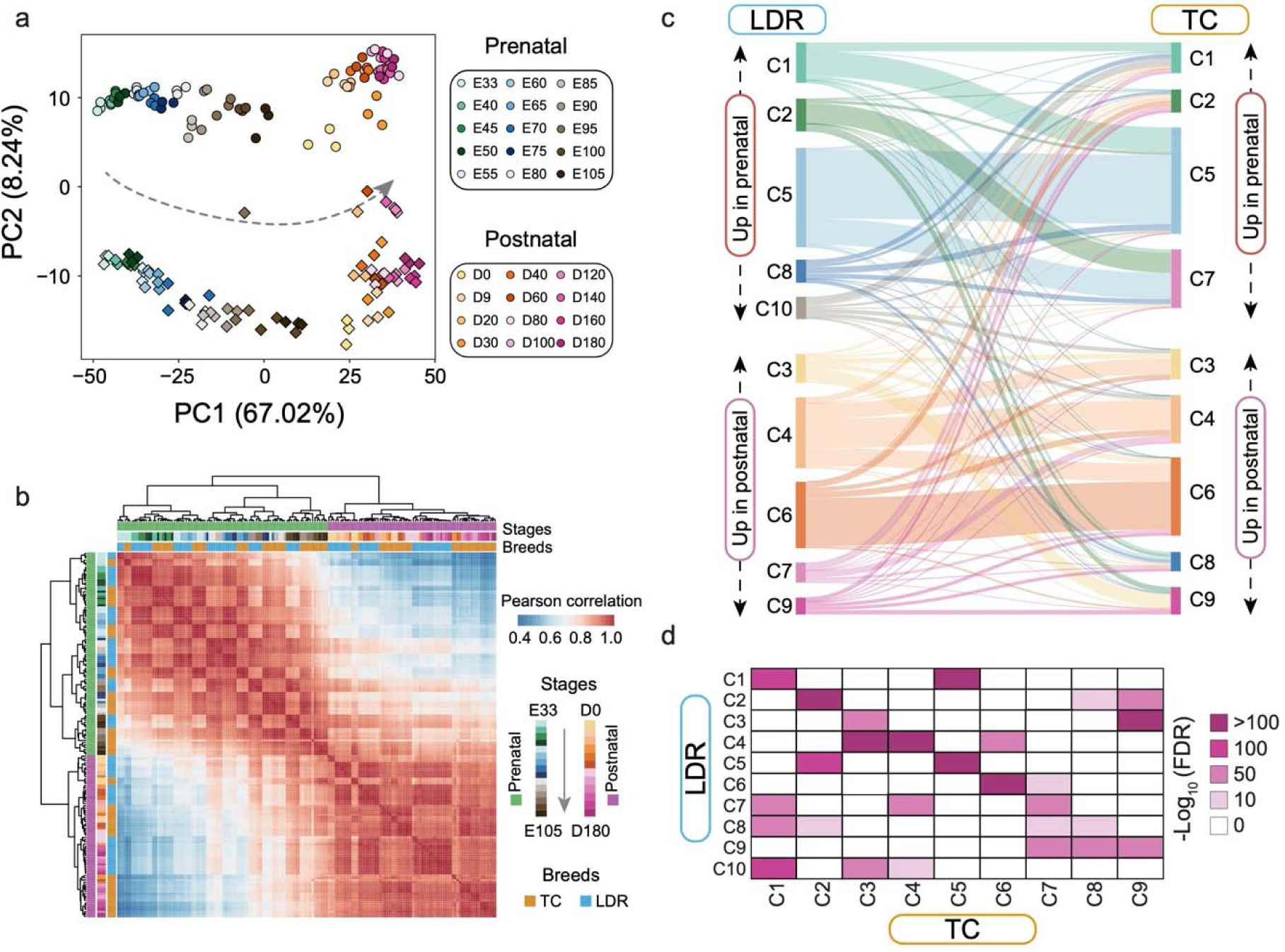
Integrative analysis of transcriptome data from Landrace (LDR) and Tongcheng (TC) pigs. **a**, PCA analysis based on the expression levels of top 1,500 variable genes from all LDR and TC individuals. A combined analysis based on all RNA-seq data from two breeds. We found a clear separation along developmental trajectories by PC1 accounting for 67.02% of the observed variance, and also robust classification of breeds by PC2 despite that this component explained only 8.24% of the variation **b**, Pearson correlation plots of all LDR and TC individuals based on gene expression. **c**, Sankey plot shows the high similarity of different clusters separately defined in LDR and TC pigs. **d**, Hypergeometric test among different clusters from LDR and TC pigs. Our results confirmed statistically significant concordance between TC and LDR, suggesting that the majority of genes coordinated myogenesis in a similar manner regardless of breeds.

**Supplementary Fig. 12.**
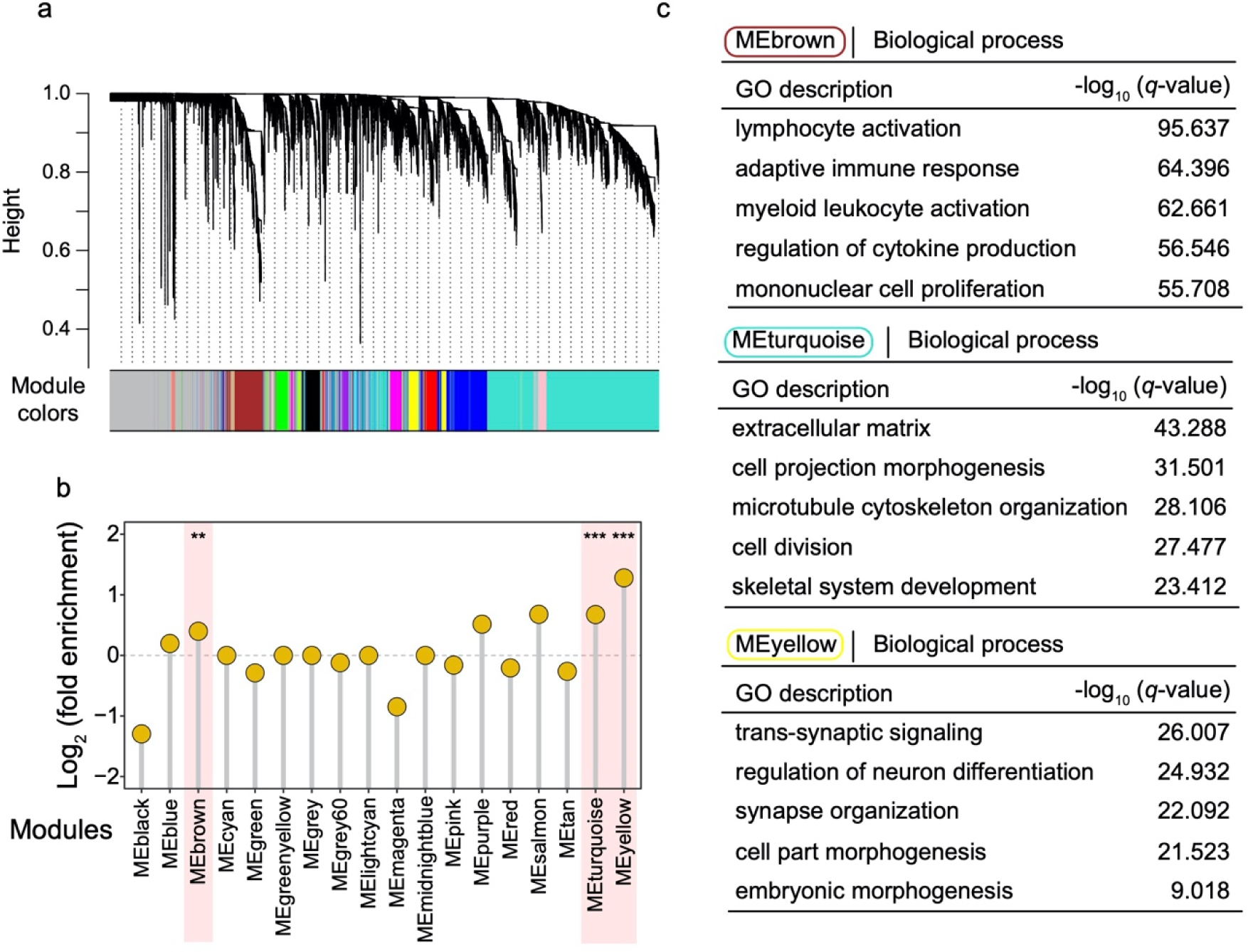
Gene expression modules affected by selective sweeps. **a**, Eighteen different modules derived by weighted gene co-expression network analysis (WGCNA) based on all LDR and TC individuals. **b**, Fold enrichment of different modules with putative swept genes. \**p* < 0.05; \*\**p* < 0.01; \*\*\**p* <0.001; ns, not significant. **c**, Enriched biological processes with the genes in MEbrown, Meturquoise and Meyellow modules.

**Supplementary Fig. 13.**
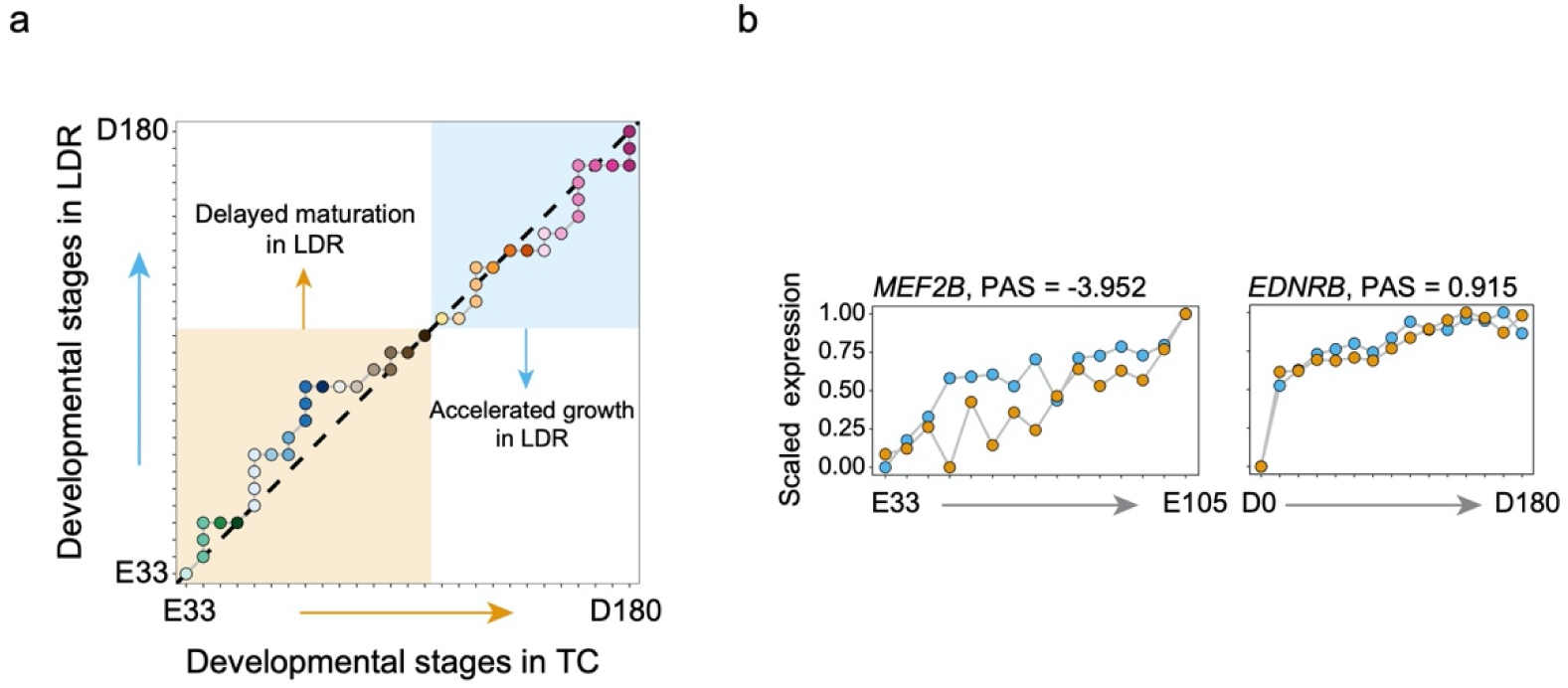
Comparison of skeletal muscle development in Landrace (LDR) and Tongcheng (TC) pigs. **a**, Developmental stage correspondences between LDR and TC pigs during myogenesis. **b**, Examples of developmental heterochrony for *MEF2B* and *EDNRB* genes.

**Supplementary Fig. 14.**
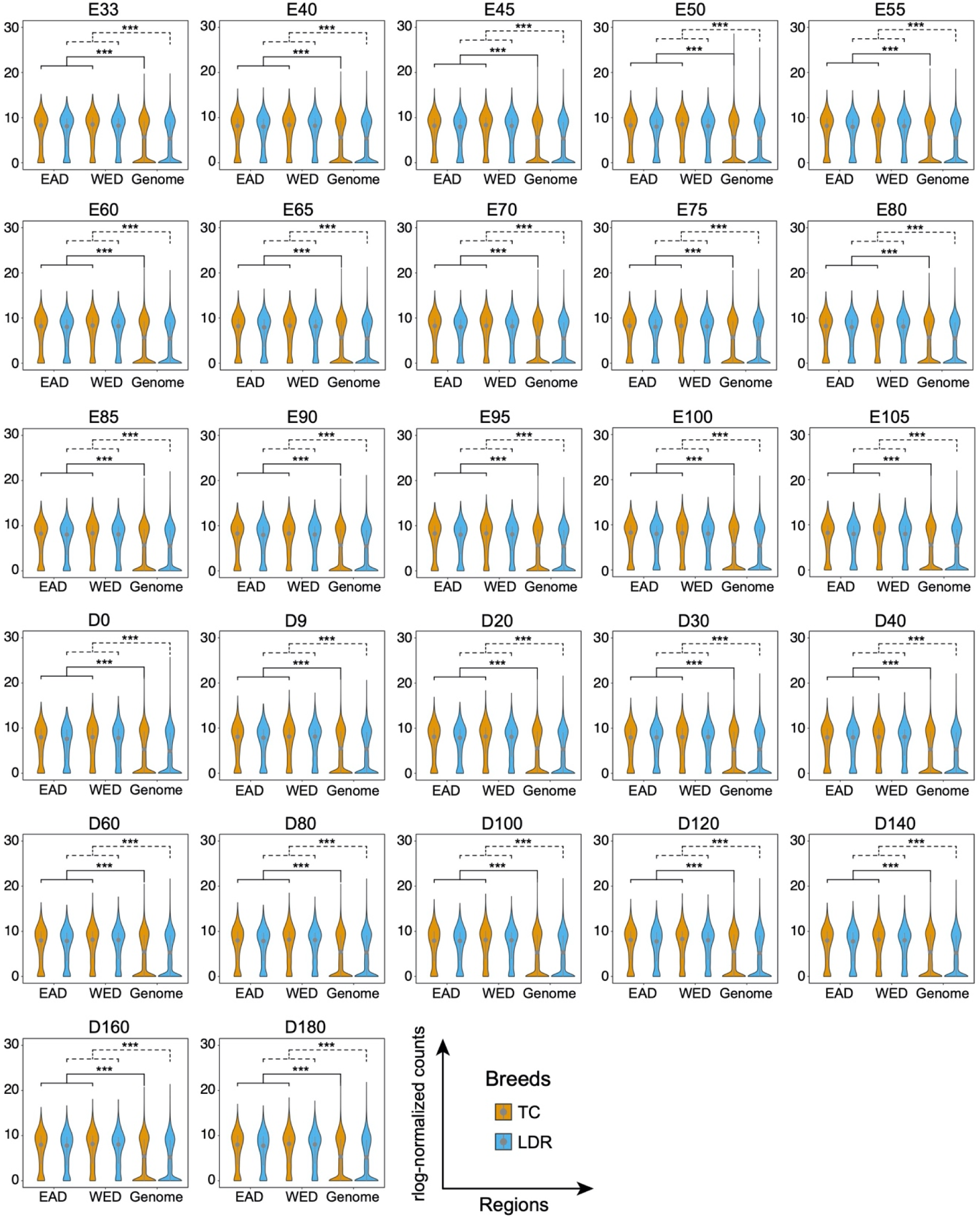
Significant differences in the expression levels between all expressed genes and the swept genes in EAD and WED groups. \*\*\**p* < 0.001.

**Supplementary Fig. 15.**
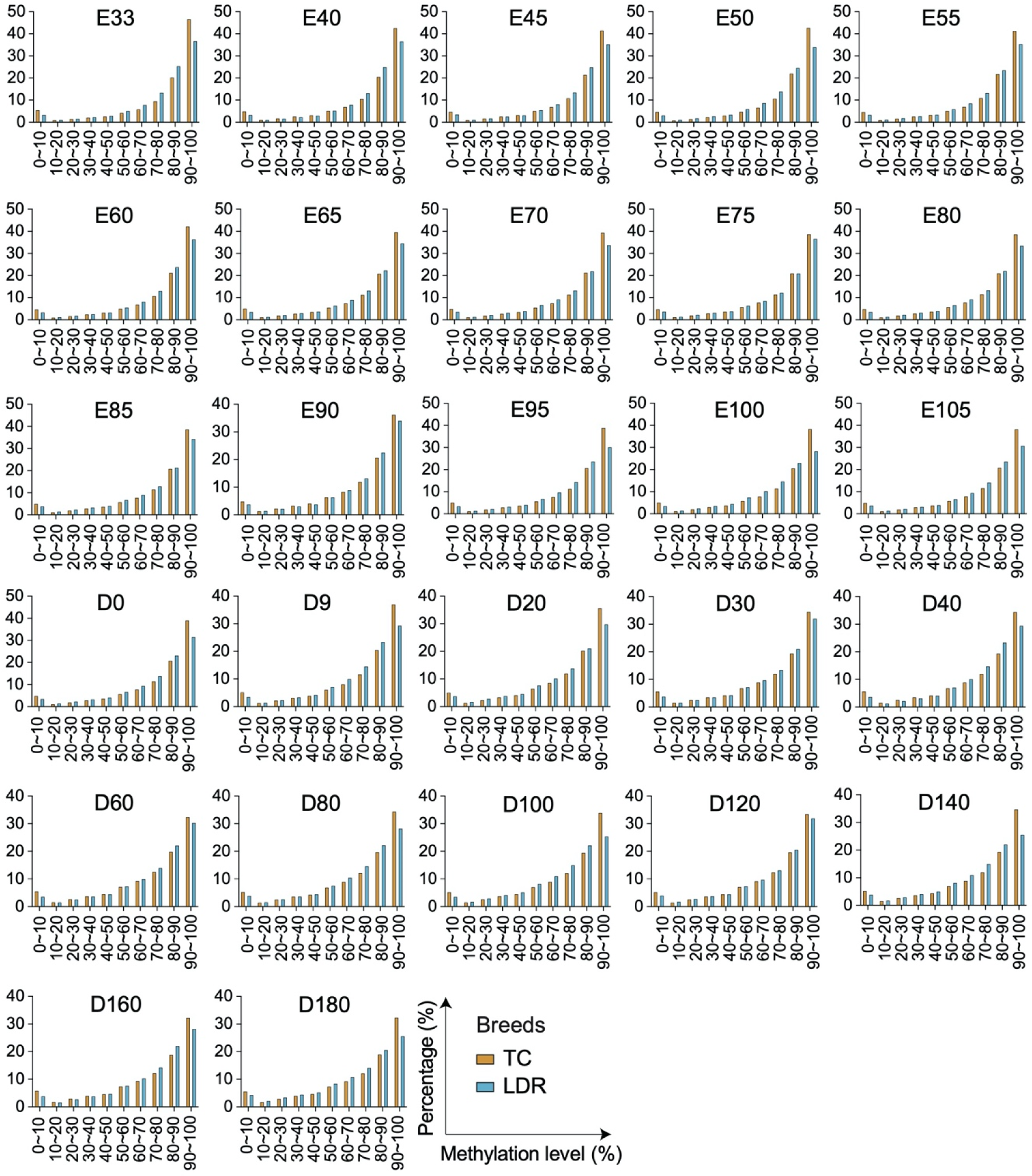
Distribution of DNA methylation profiles across the 27 skeletal muscle developmental stages in Landrace (LDR) and Tongcheng (TC) pigs. The DNA methylation levels are divided evenly into ten groups. We found an average of 72.61% and 71.63% methylation over all CpG sites in TC and LDR genomes respectively, and the majority of sites at each developmental stage showed very high methylation states of greater than 80%.

**Supplementary Fig. 16.**
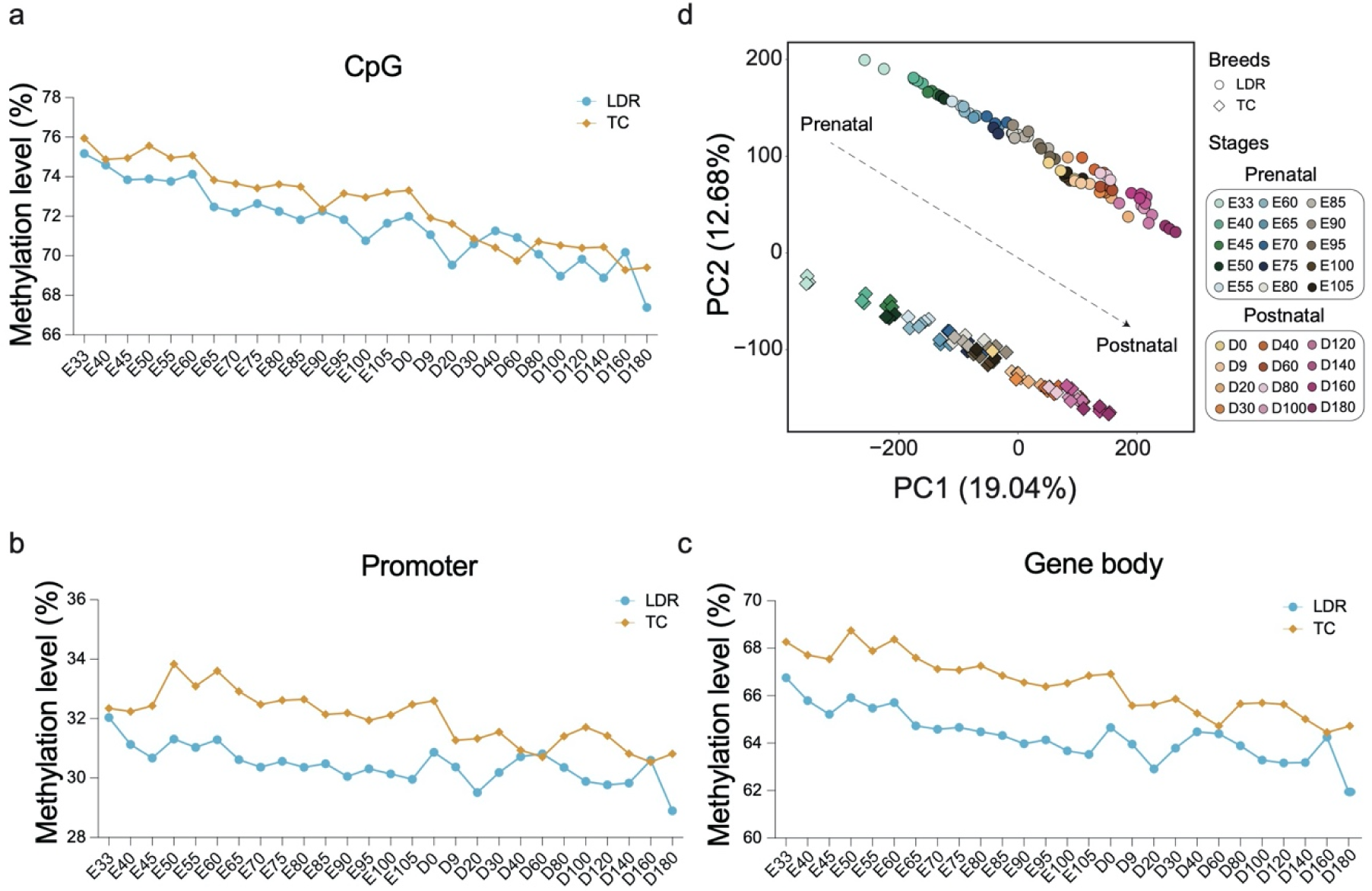
Comparisons of DNA methylation levels between Landrace (LDR) and Tongcheng (TC) pigs. **a-c**, Dynamic DNA methylation landscape for al CpG sites (a), gene body (b) and promoter regions (c) in LDR and TC breeds. The gradual decrease of methylation levels in all CpG sites were detected throughout the whole development, with the biggest difference close to 10% from prenatal day 33 (E33) to postnatal day 180 (D180), which were similar between TC and LDR. Notably, the TC breed exhibited much heavier CpG methylation levels than LDR across the whole genome in almost all 27 developmental stages, and the same results were found in the promoters and gene bodies. **d**, PCA analysis based on the DNA methylation levels of LDR and TC pigs.

**Supplementary Fig. 17.**
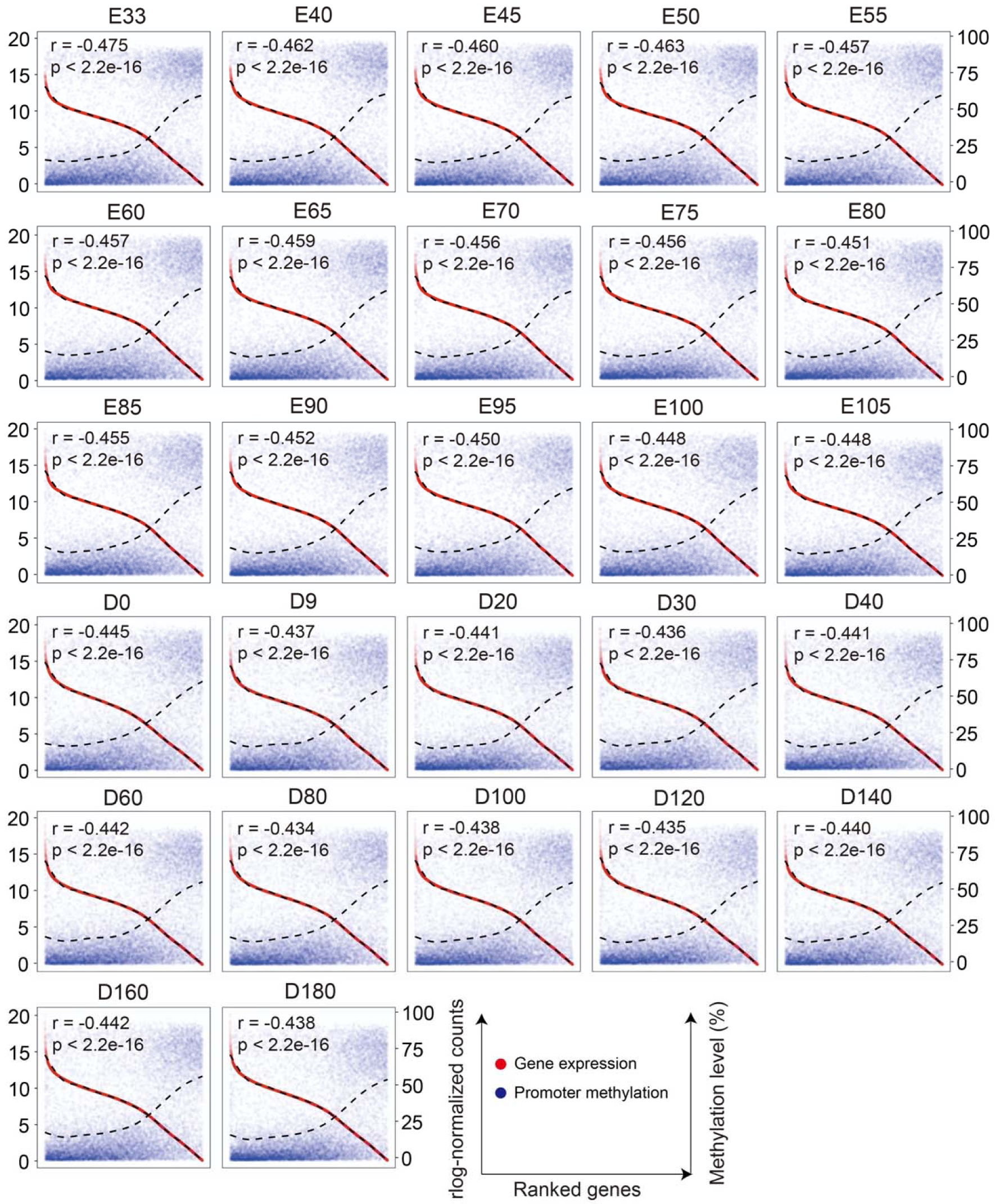
Correlation between gene expression and DNA methylation levels of promoter regions in Landrace pig. The Pearson correlation coefficients (*r*) between gene expression levels and DNA methylation levels were calculated. The dotted fitting curves separately represent gene expression levels and DNA methylation levels. The horizontal axis from left to right below each box represents the expression gene levels from high to low. The promoter-associated CpG islands have profound suppression effects in transcriptional levels with negative correlation coefficients close to -0.500.

**Supplementary Fig. 18.**
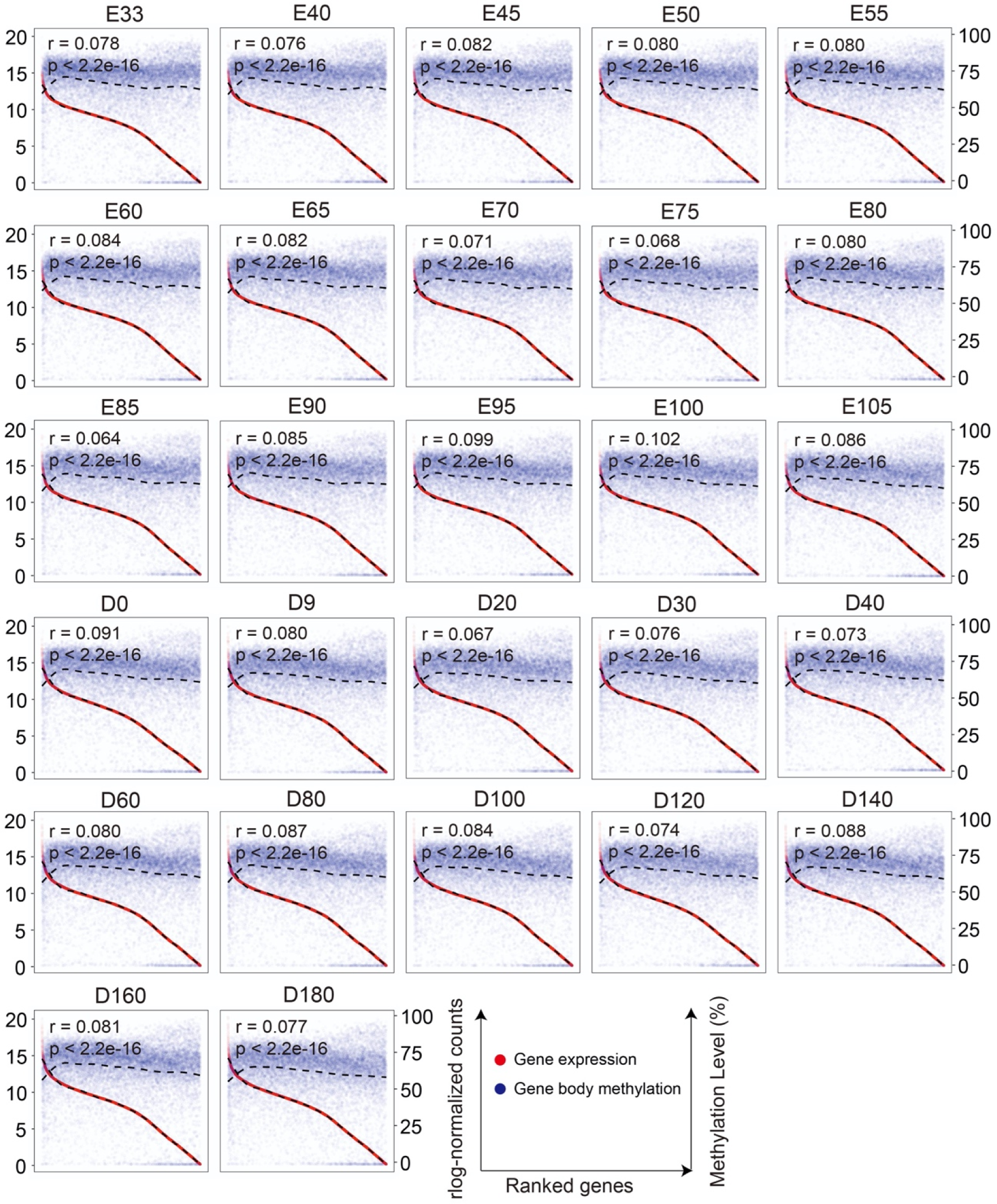
Correlation between gene expression and DNA methylation levels of gene body regions in Landrace pig. The Pearson correlation coefficients (*r*) between gene expression levels and DNA methylation levels were calculated. The dotted fitting curves separately represent gene expression levels and DNA methylation levels. The horizontal axis from left to right below each box represents the gene expression levels from high to low. Only weak correlations between DNA methylation and gene expression were found across the gene body regions, suggesting a more complex regulatory manner of gene body DNA methylation.

**Supplementary Fig. 19.**
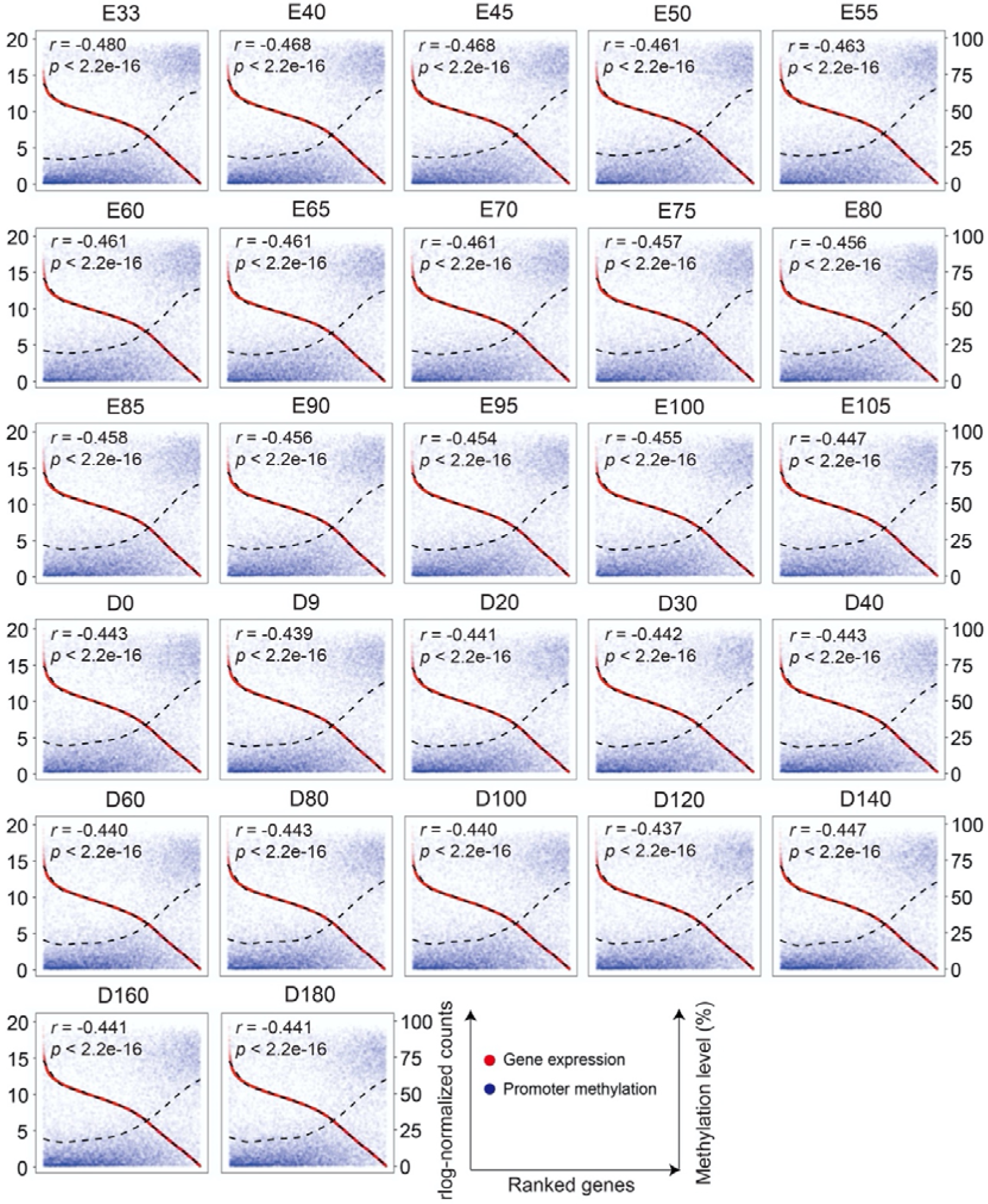
Correlation between gene expression and DNA methylation levels of promoter regions in Tongcheng pig. The Pearson correlation coefficients (*r*) between gene expression levels and DNA methylation levels were calculated. The dotted fitting curves separately represent gene expression levels and DNA methylation levels. The horizontal axis from left to right below each box represents the gene expression levels from high to low.

**Supplementary Fig. 20.**
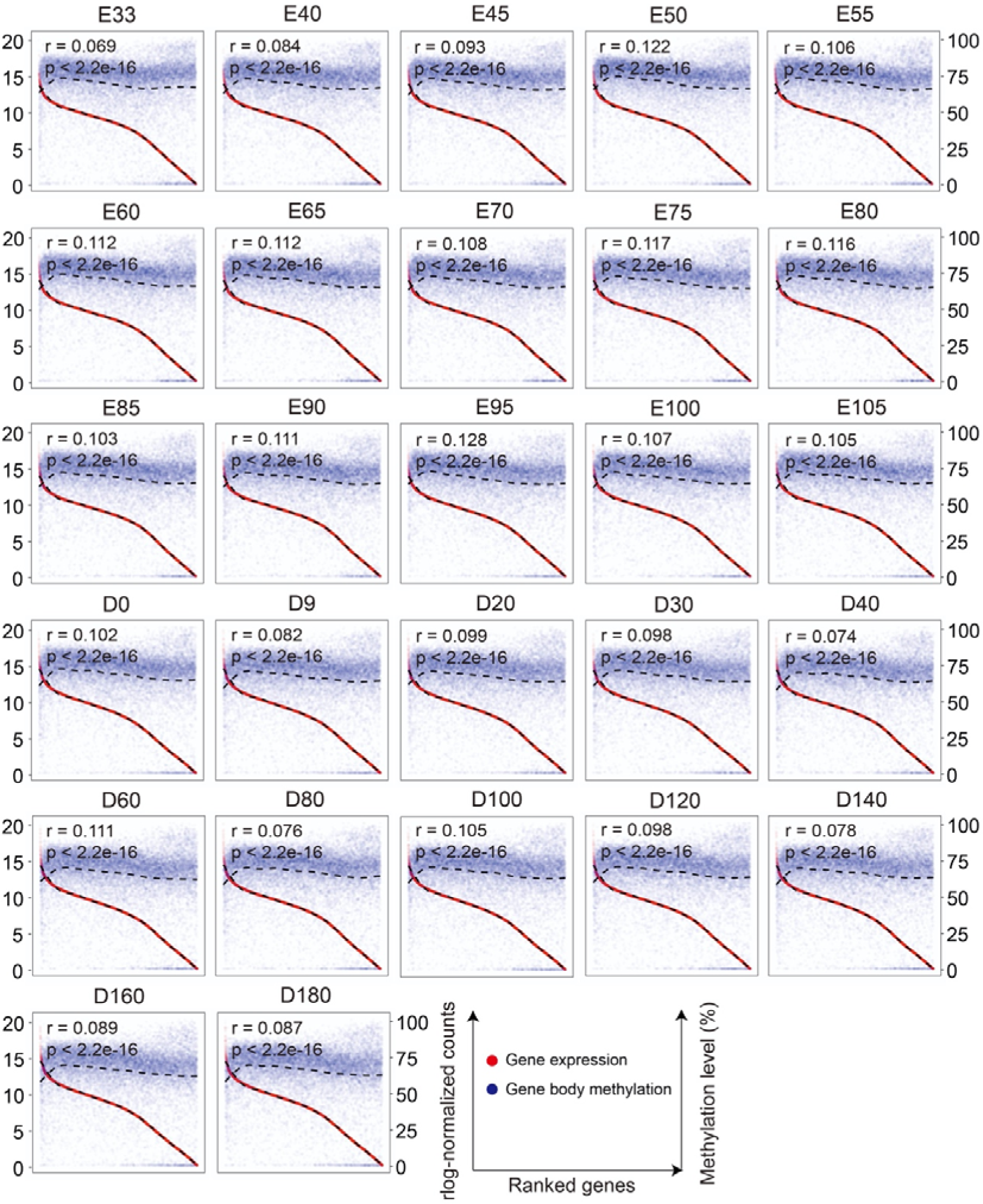
Correlation between gene expression and DNA methylation levels of gene body regions in Tongcheng pig. The Pearson correlation coefficients (*r*) between gene expression levels and DNA methylation levels were calculated. The dotted fitting curves separately represent gene expression level and DNA methylation level. The horizontal axis from left to right below each box represents the gene expression levels from high to low.

**Supplementary Fig. 21.**
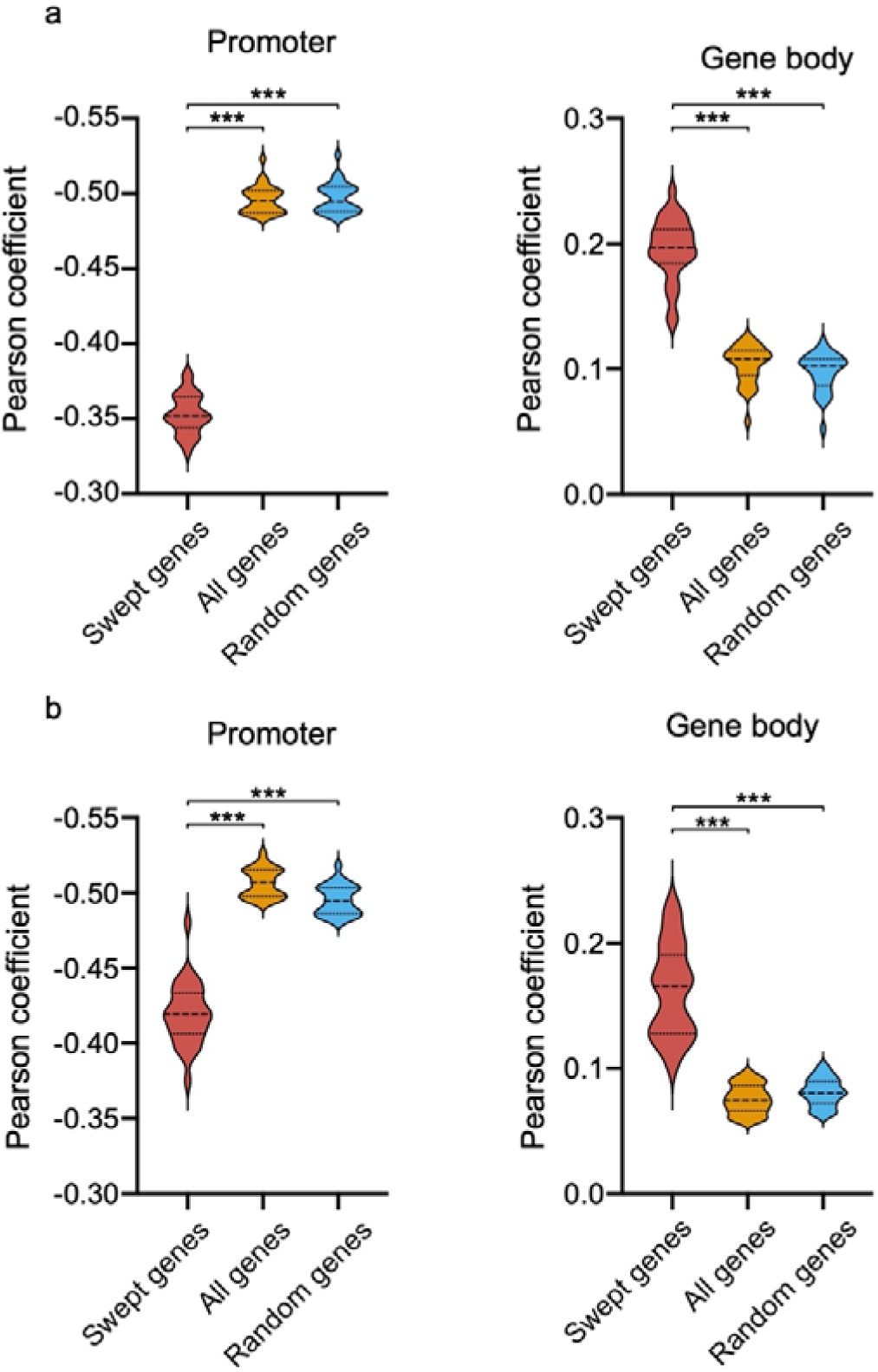
Changed correlation patterns between gene expression and DNA methylation levels by selective sweeps. **a-b**, Correlation coefficients of swept genes are lower for promoter regions, but higher for gene body regions in both LDR (a) and TC (b) pigs, compared with these of all genes and randomly selected genes. The number of random genes was matched to swept genes in EAD and WED groups, and the Pearson correlation coefficients are calculated based on 1,000 permutations. \*\*\**p* < 0.001.

**Supplementary Fig. 22.**
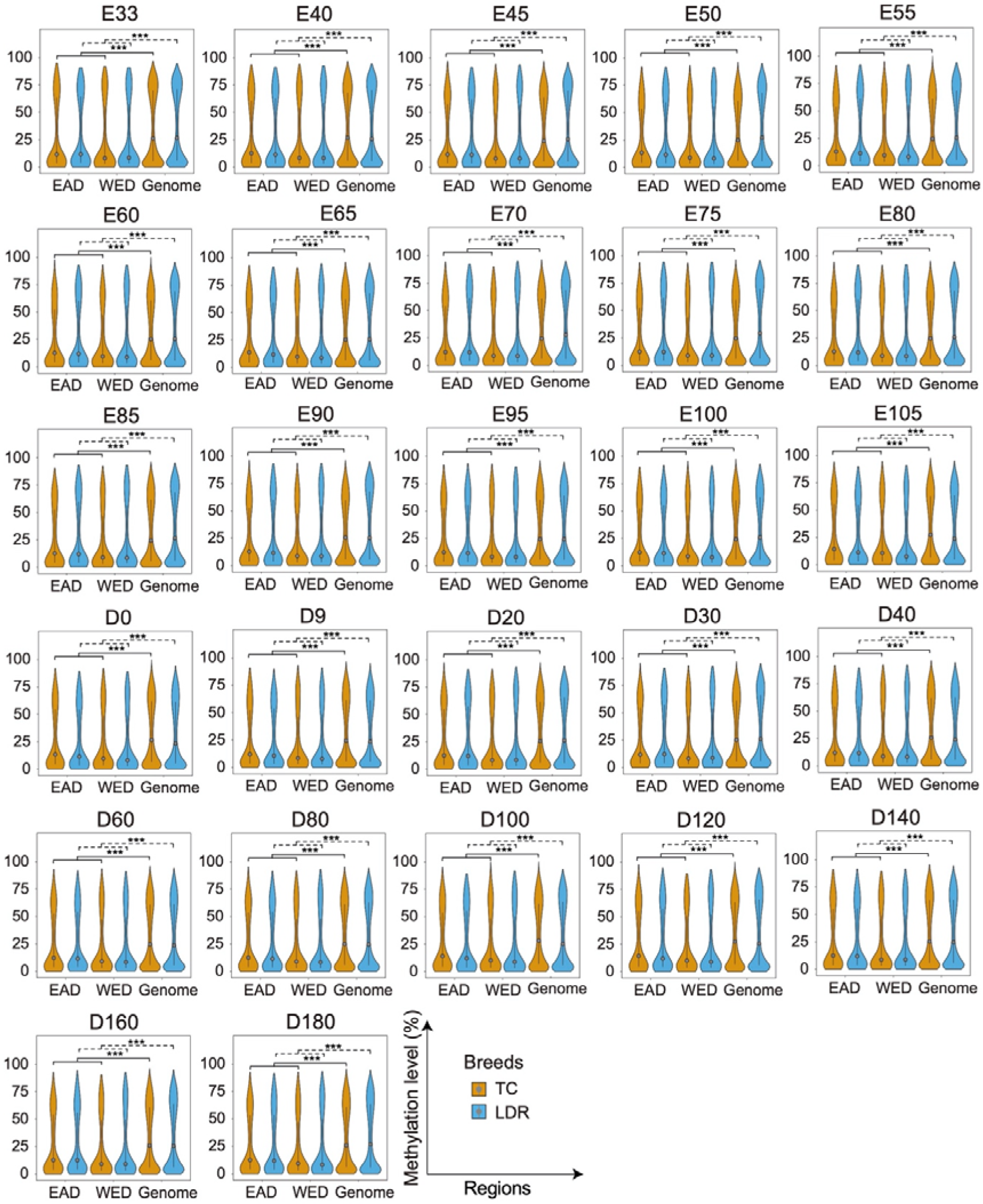
Comparison of the DNA methylation levels in promoter regions for all genes and swept genes. \*\*\**p* < 0.001.

**Supplementary Fig. 23.**
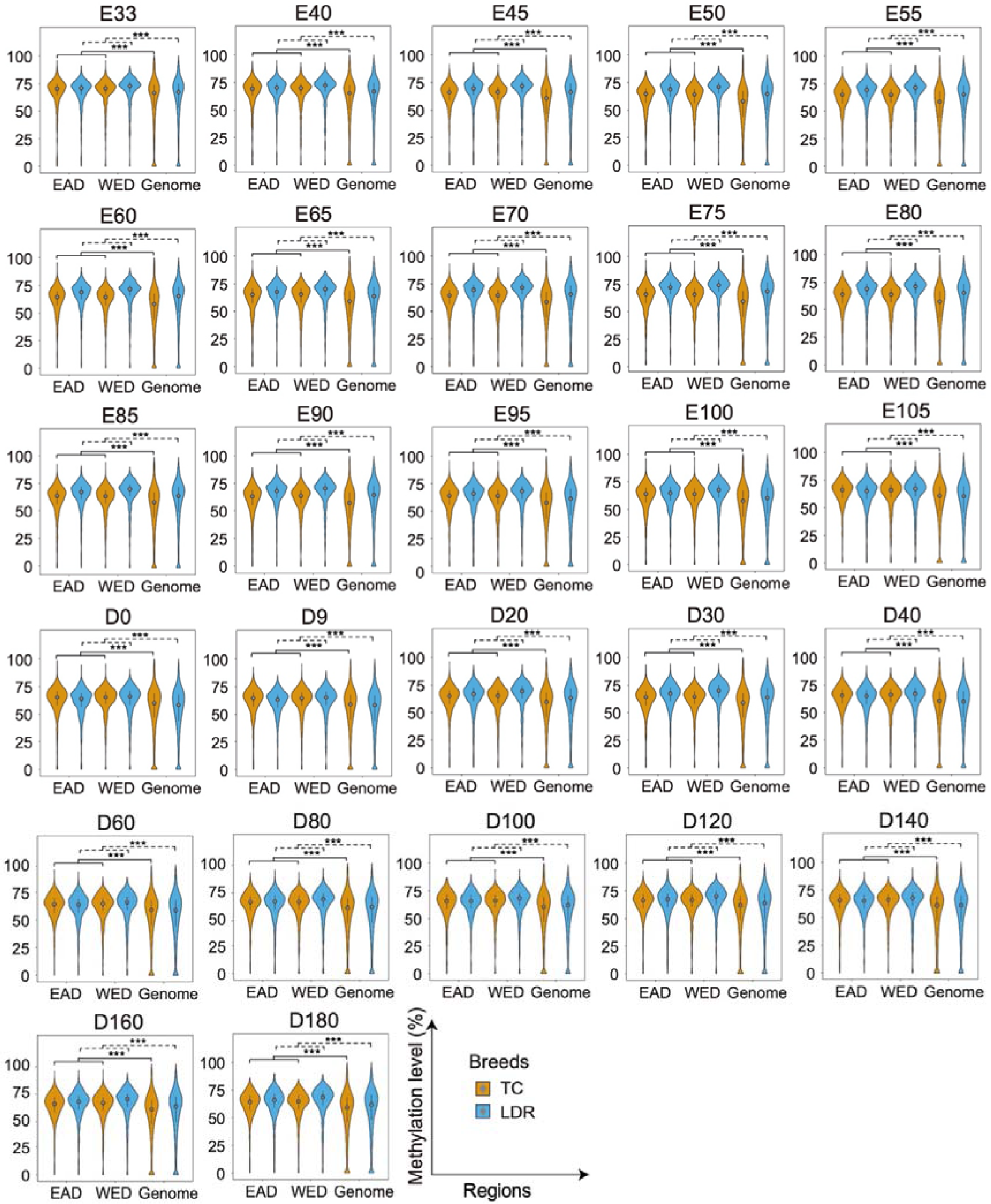
Comparison of the DNA methylation levels in gene body regions for all genes and swept genes. \*\*\**p* < 0.001.

**Supplementary Fig. 24.**
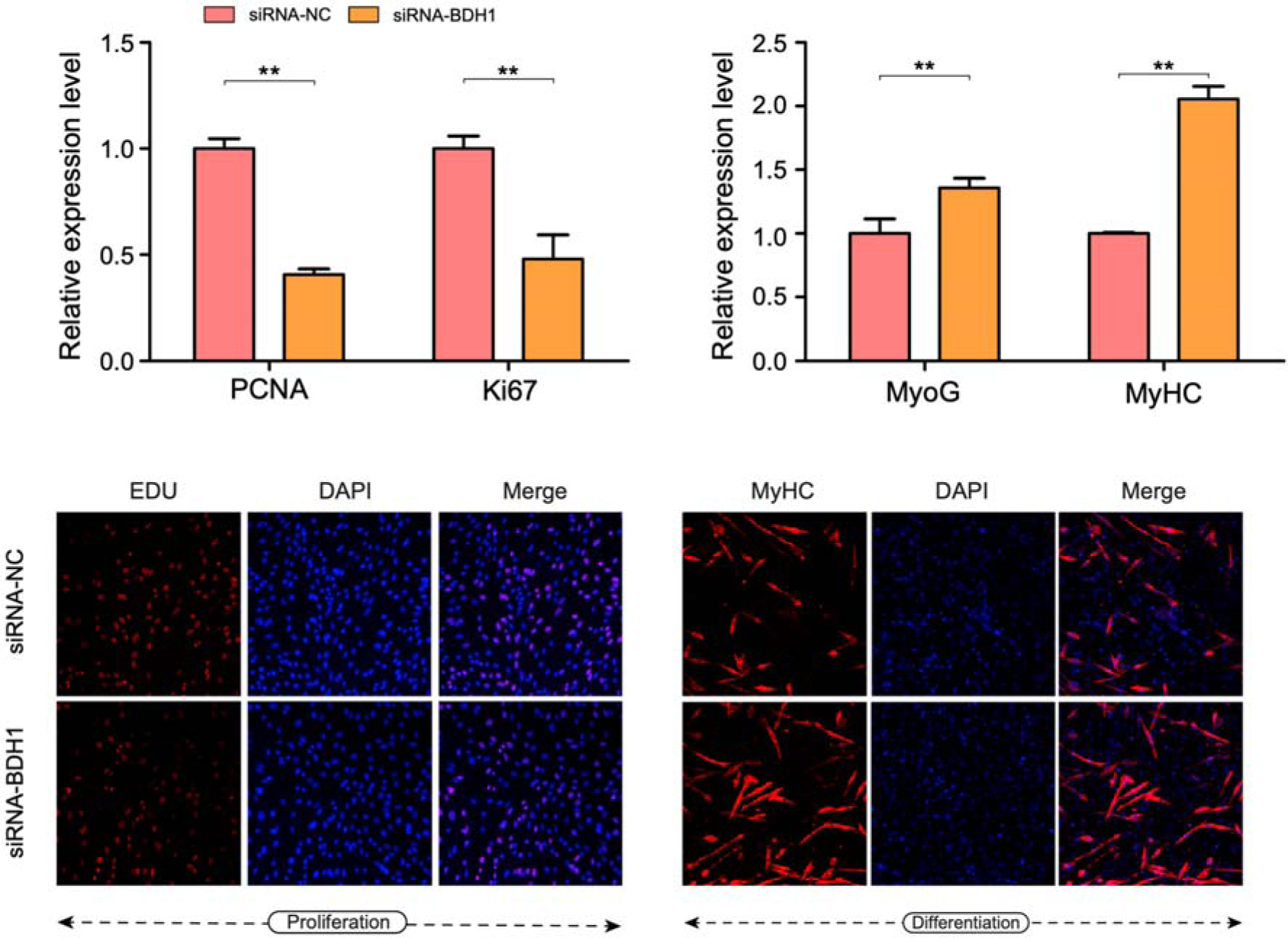
Cell differentiation assessment for the expression levels of *PCNA*, *Ki67*, *MyoG*, and *MyHC* marker upon *BDH1* knockdown by qRT-PCR analysis in C2C12 cells.

**Supplementary Fig. 25.**
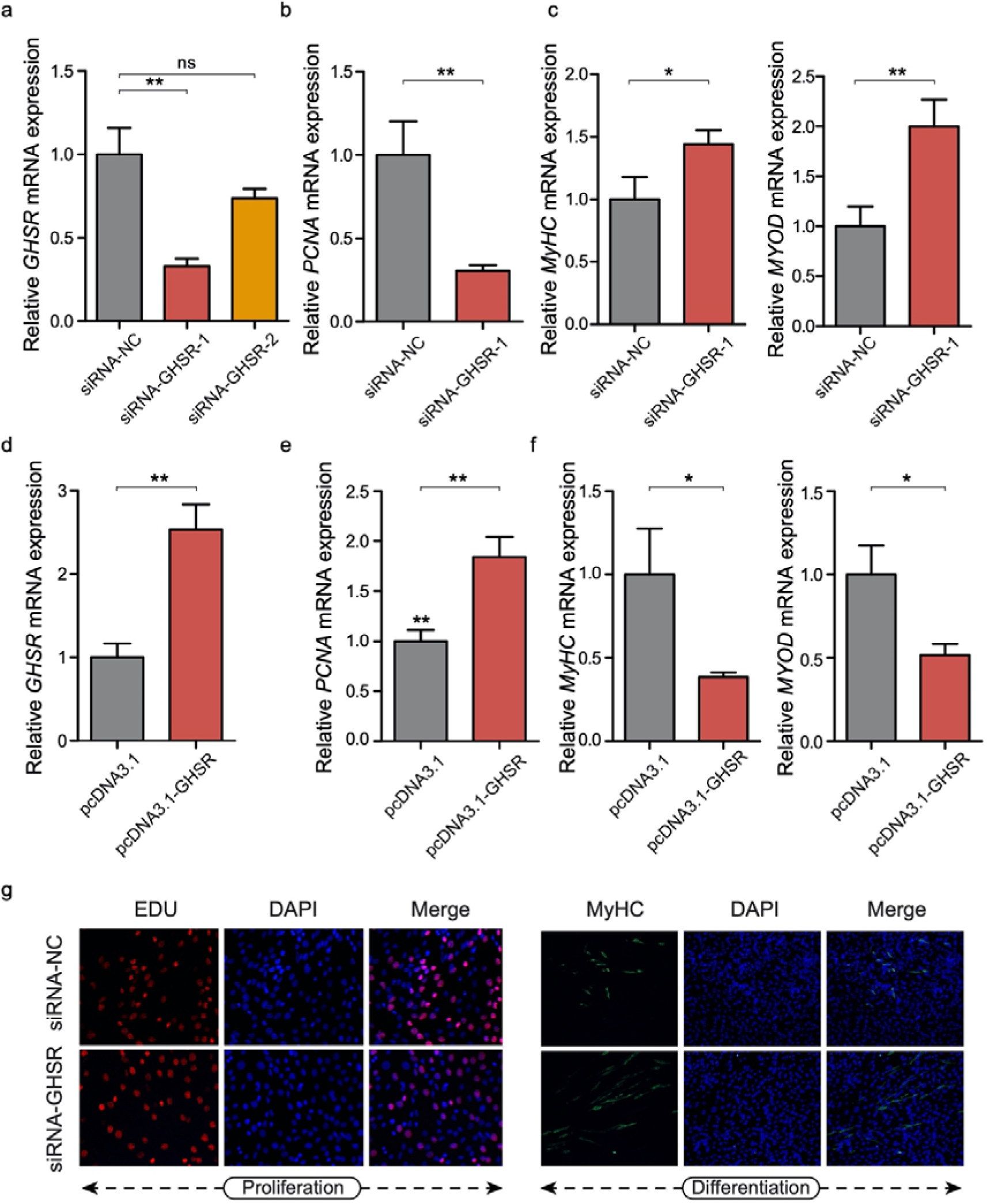
Enhanced capacity for cell proliferation of *GHSR* gene. **a**, Knockdown efficiency of *GHSR* in C2C12 cells. \**p* < 0.05; \*\**p* < 0.01; \*\*\**p* < 0.001; ns, not significant. **b**, Cell proliferation assessment for the expression level of *PCNA* marker upon *GHSR* knockdown by qRT-PCR analysis in C2C12 cells. **c**, Cell differentiation assessment for the expression levels of *MyHC* and *MYOD* marker upon *GHSR* knockdown by qRT-PCR analysis in C2C12 cells. **d**, Overexpression efficiency of *GHSR* in C2C12 cells. **e**, Cell proliferation assessment for the expression level of *PCNA* marker upon *GHSR* overexpression by qRT-PCR analysis in C2C12 cells. **f**, Cell differentiation assessment for the expression levels of *MyHC* and *MYOD* marker upon *GHSR* overexpression by qRT-PCR analysis in C2C12 cells. **g**, Cell proliferation and differentiation experiments measured by EDU and MyHC immunofluorescence upon *GHSR* knockdown.

**Supplementary Fig. 26.**
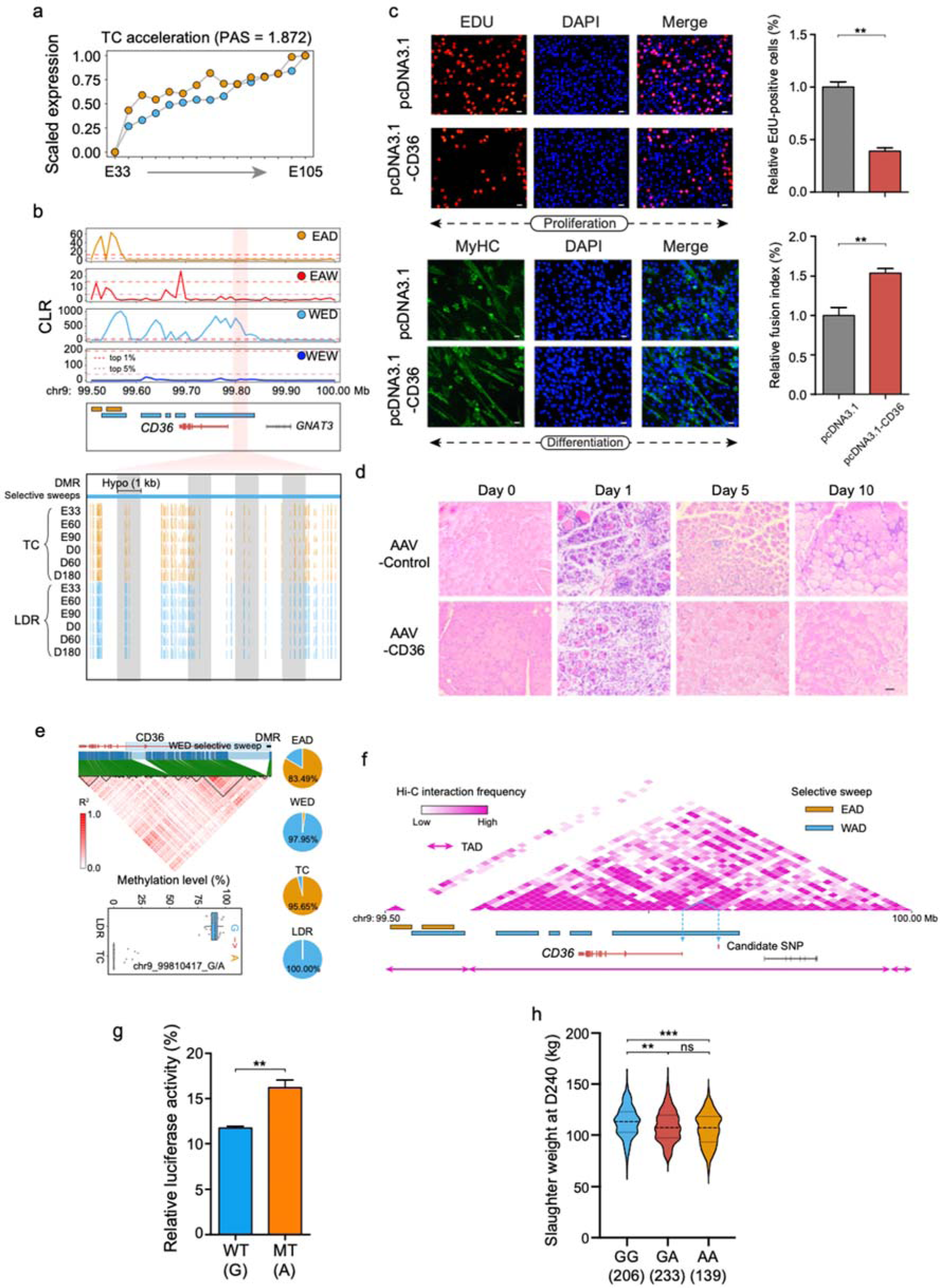
Comprehensive analysis of the *CD36* gene associated with skeletal muscle development and meat performance. **a**, More advanced progression of *CD36* at the TC prenatal stage. **b**, Visualization of methylation tracks around the swept *CD36* region. Dashed purple and red lines represent thresholds of the top 5% and 1% CLR values in the *CD36* region, respectively. **c**, Cell proliferation and differentiation assays measured by EDU and MyHC immunofluorescence upon CD36 overexpression. **d**, H&E staining for regenerating muscle on days 0, 1, 5 and 10 after cardiotoxin (CTX) injury in AAV-mediated non-target control and pcDNA3.1-CD36 groups, respectively. Representative images are shown at 20× magnification (scale bars = 100 μm). **e**, Distinct methylation patterns and genotype spectrum between two alleles in the CpG-SNP site showing strong LD with the *CD36* gene. **f**, The candidate SNP and the CD36 gene located in the same topologically associated domain (TAD) defined by Hi-C data. **g**, Enhancer activities of two alleles by the luciferase reporter assay in the HEK293T cells. **h**, Significant phenotypic difference in slaughter weight among three genotypes.

**Supplementary Fig. 27.**
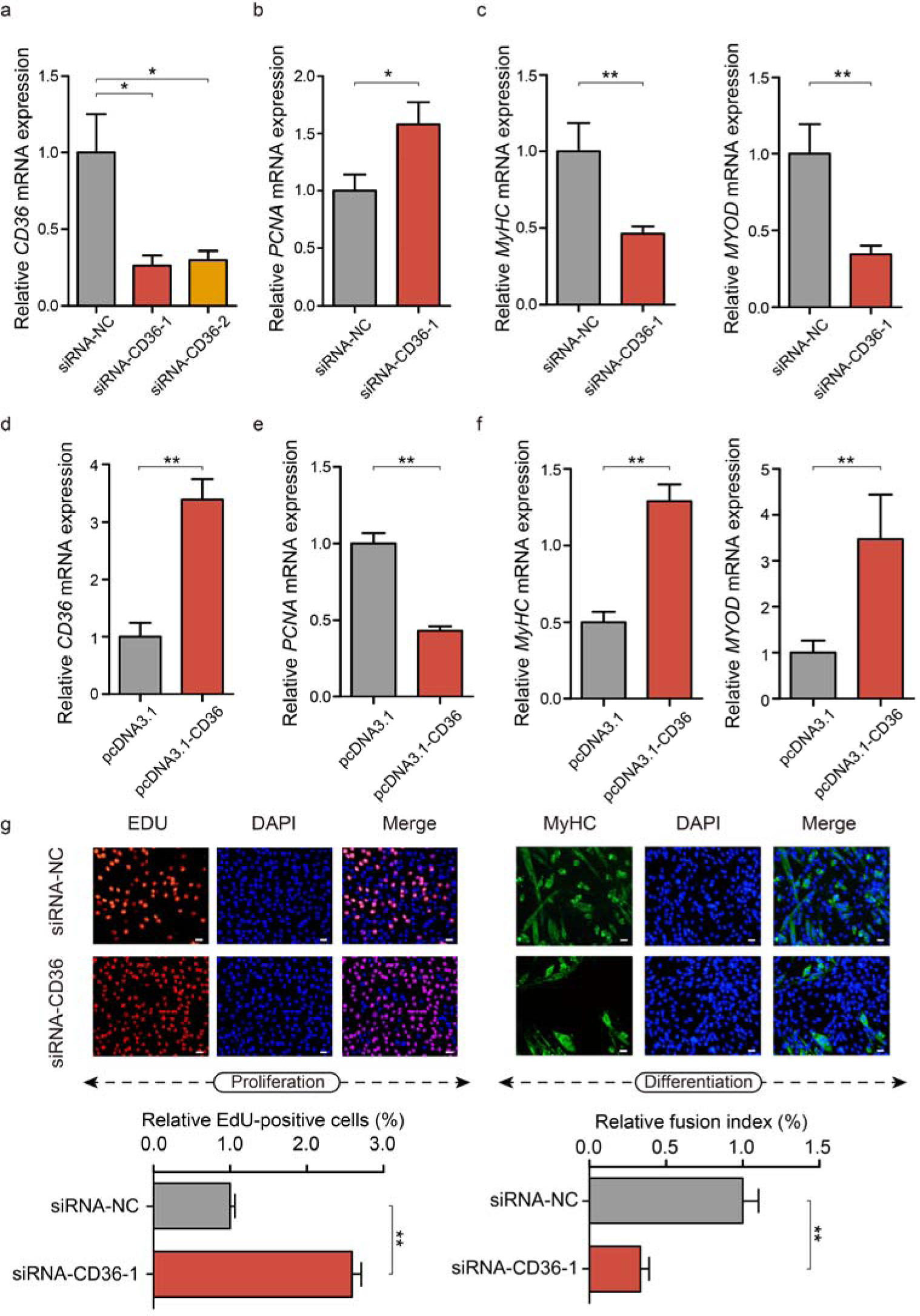
Enhanced capacity for cell differentiation of *CD36* gene. **a**, Knockdown efficiency of *CD36* in C2C12 cells. \**p* < 0.05; \*\**p* < 0.01; \*\*\**p* < 0.001; ns, not significant. **b**, Cell proliferation assessment for the expression level of *PCNA* marker upon *CD36* knockdown by qRT-PCR analysis in C2C12 cells. **c**, Cell differentiation assessment for the expression levels of *MyHC* and *MYOD* marker upon *CD36* knockdown by qRT-PCR analysis in C2C12 cells. **d**, Overexpression efficiency of *CD36* in C2C12 cells. \*\*\**p* < 0.001. **e**, Cell proliferation assessment for the expression level of *PCNA* marker upon *CD36* overexpression by qRT-PCR analysis in C2C12 cells. **f**, Cell differentiation assessment for the expression levels of *MyHC* and *MYOD* marker upon *CD36* overexpression by qRT-PCR analysis in C2C12 cells. **g**, Cell proliferation and differentiation experiments measured by EDU and MyHC immunofluorescence upon *CD36* knockdown.

**Supplementary Fig. 28.**
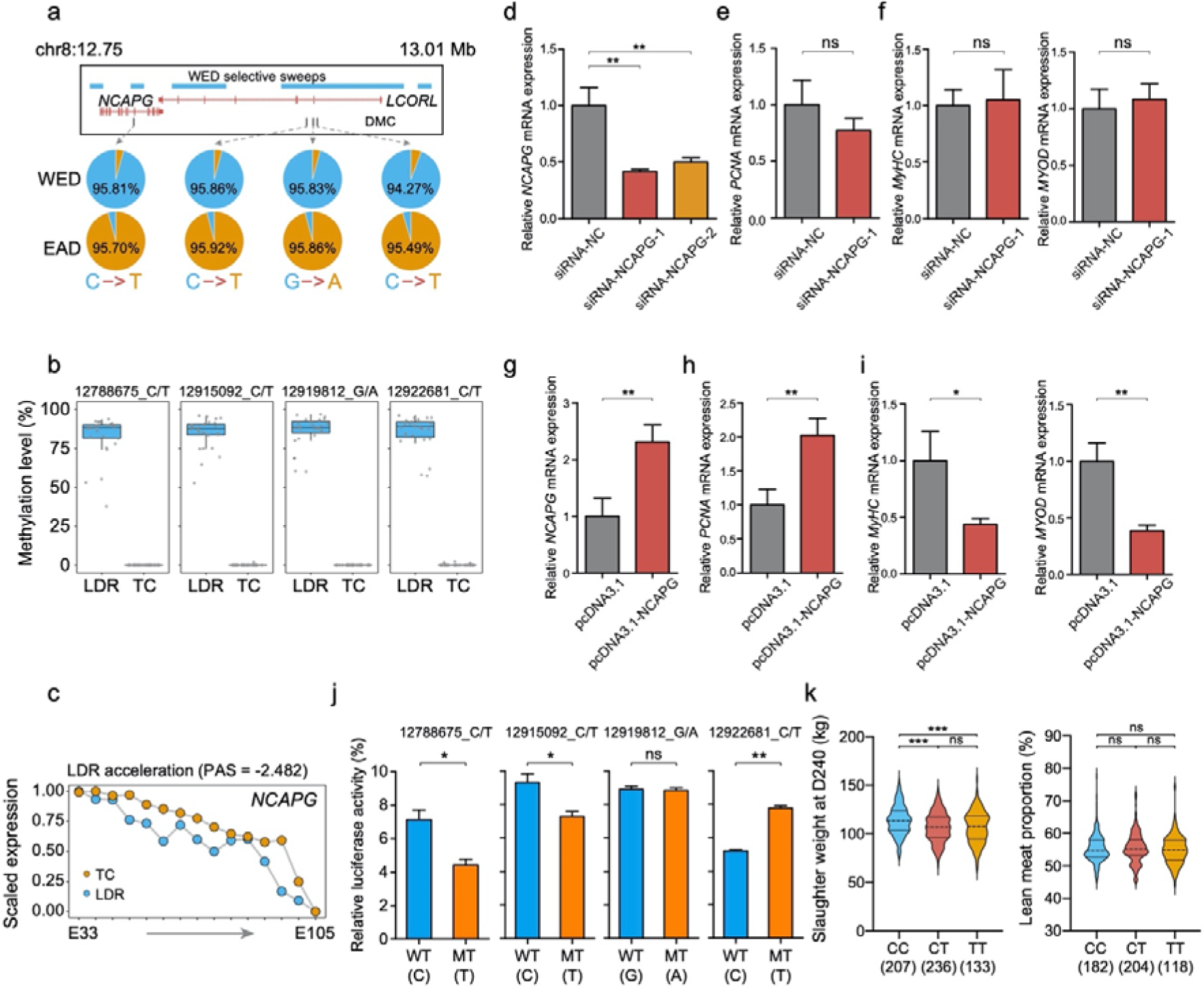
Comprehensive functional analysis of *NCAPG*-*LCORL* locus. **a**, Four fixed SNPs from methylated CpG to TpG/CpA in the *NCAPG*-*LCORL* locus. **b**, Methylation levels of the four CpG-SNPs in LDR and TC pigs. **c**, Temporal advanced expression pattern of *NCAPG* gene in the LDR breed. **d**, Knockdown efficiency of *NCAPG* in C2C12 cells. \**p* < 0.05; \*\**p* < 0.01; \*\*\**p* < 0.001; ns, not significant. **e**, Cell proliferation assessment for the expression level of *PCNA* marker upon *NCAPG* knockdown by qRT-PCR analysis in C2C12 cells. **f**, Cell differentiation assessment for the expression levels of *MyHC* and *MYOD* marker upon *NCAPG* knockdown by qRT-PCR analysis in C2C12 cells. **g**, Overexpression efficiency of *NCAPG* in C2C12 cells. **h**, Cell proliferation assessment for the expression level of *PCNA* marker upon *NCAPG* overexpression by qRT-PCR analysis in C2C12 cells. **i**, Cell differentiation assessment for the expression levels of *MyHC* and *MYOD* marker upon *NCAPG* overexpression by qRT-PCR analysis in C2C12 cells. **j**, Comparisons of the effects of the four differential DNA variants on enhancer activity by luciferase reporter assays in HEK293T cells. **k**, Phenotypic consequences of the intronic SNP in the *NCAPG* in the slaughter weight at day 240 and lean meat proportion.

